# Gene expression in isolated cowpea (Vigna unguiculata L. Walp) cells from meiosis to seed initiation

**DOI:** 10.1101/2020.01.17.909945

**Authors:** Nial Gursanscky, Danielle Mazurkiewicz, Martina Juranić, Susan D. Johnson, Gloria León, Rocio Escobar-Guzmán, Rigel Salinas-Gamboa, Itzel Amasende-Morales, Matteo Riboni, Melanie Hand, Andrew Spriggs, Jean-Philippe Vielle-Calzada, Anna M.G. Koltunow

## Abstract

Molecular knowledge of pathways regulating seed formation in legumes, remains scarce. Thirteen isolated cell-type transcriptomes were developed, spanning temporal events of male and female gametogenesis and seed initiation, to examine pathways involved in cowpea seed formation. *In situ* hybridization confirmed localization of *in silico* identified cell-specific genes, verifying transcriptome utility. Cowpea and *Arabidopsis* reproductive cells showed some conservation in regulators enabling cell-type expression as some cowpea cell-specific genes promoters and their *Arabidopsis* homologs directed expression to identical reproductive cell-types in transgenic plants. *In silico* analyses revealed gene expression similarities and differences with genes in pathways regulating reproductive events in other plants. Meiosis-related genes were expressed at mitotic stages of gametogenesis and during sporophytic development in cowpea. Plant hormone pathways showing preferential expression at particular reproductive stages were identified. Expression of epigenetic pathways, resembling those found in *Arabidopsis,* including microRNA mediated gene silencing, RNA directed DNA methylation and histone modification were associated with particular stages of male and female gametophyte development, suggesting roles in gametogenic cell specification and elaboration. Analyses of cell-cycle related gene expression in mature cowpea female gametophytes, indicated that the egg and central cell were arrested at the G1/S and G2/M cell cycle phases, respectively, prior to fertilization. Pre-fertilization female gametophyte arrest was characterized by barely detectable auxin biosynthesis gene expression levels, and elevated expression of genes involved in RNA-mediated gene silencing and histone modification. These transcriptomes provide a useful resource for additional interrogation to support functional analyses for development of higher yielding cowpea and syntenic legume crops.

**One sentence summary:** Analyses of laser capture derived cell-type transcriptomes spanning meiosis to seed initiation revealed gene expression profiles during cell specification and reproductive development in cowpea.

## Introduction

Legumes contribute directly and indirectly to world food supply. Higher in protein, they provide forage for animals and increase soil fertility by fixing nitrogen. Agronomic improvement of eight grain legumes, including cowpea (*Vigna unguiculata* L. Walp) has been targeted by the Consultative Group for International Agricultural Research (https://storage.googleapis.com/cgiarorg/2018/11/SC7-B_Breeding-Initiative-1.pdf). We aim to increase seed yield and quality in cowpea, which originated in Africa and remains an important subsistence crop in sub-Saharan Africa (Singh, 2014). Although cowpea is relatively tolerant to drought, it is prone to high levels of flower and pod drop, resulting in decreased seed yields. There is limited knowledge of the molecular pathways supporting successful elaboration of cowpea male and female gametophytes and seeds.

Current genomic resources for cowpea include a reference genome (Lonardi et al., 2019, Munoz-Amatriain et al., 2017) together with survey genomes and transcriptomes from various plant tissues (Spriggs et al., 2018, Yao et al., 2016). Reproductive events occur only in a few cell-types buried within cowpea floral organs (Salinas-Gamboa et al., 2016). Therefore, laser capture microdissection (LCM) is a useful approach to isolate reproductive cell-types to generate informative transcriptomes. Interrogation of these reproductive cell-type transcriptomes in conjunction with cowpea genome and tissue transcriptome resources could accelerate identification and functional examination of pathways supporting successful cowpea seed formation.

A cowpea reproductive calendar has been developed linking morphological floral features in cowpea to reproductive events in floral tissues to support collection of tissues for molecular analyses (Salinas-Gamboa et al., 2016). Male gametophyte precursor cells, termed pollen mother cells (PMC) undergo meiosis (sporogenesis) forming tetrads of microspores (mTET) in the anther. Individual microspores (MIC) then undergo mitosis (microgametogenesis) and differentation to form the mature pollen grain (MP-SC) containing a generative and a vegetative cell (Fig. 1A). By contrast, during female gametophyte development, a single megaspore mother cell (MMC) in each ovule undergoes meiosis to form a tetrad of megaspores (fTET). Three megaspores degenerate and the surviving functional megaspore (FM) undergoes three rounds of mitosis to form a syncytium of eight nuclei (megagametogenesis). Cellularization events result in a 7-celled *Polygonum*-type female gametophyte, or embryo sac, and three of these degrade. The mature cowpea embryo sac (MES) contains an egg cell (EC) progenitor of the embryo (Em), flanked by two synergids (Sy), which guide fertilization events and the central cell (CC) containing two fused haploid nuceli, which is the progenitor of the endosperm (En; Fig. 1A; Salinas-Gamboa et al., 2016).

**Figure 1.**
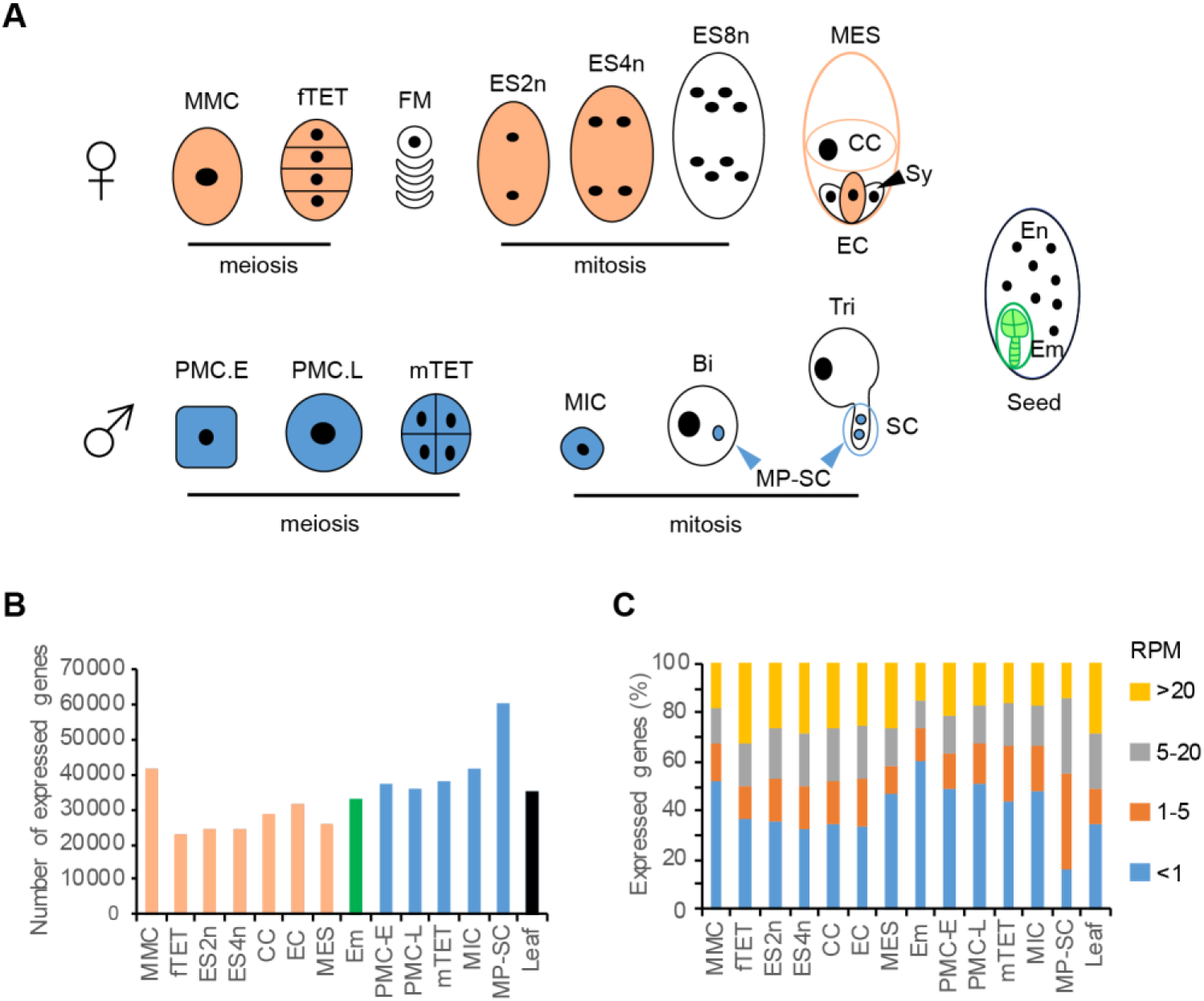
Stages of reproductive development in cowpea and gene expression in developed transcriptomes. A, Cartoon of temporal stages of male and female gametophyte development showing cell-type stages collected by LCM for RNAseq. For detail refer to text. B, The number of genes expressed in each transcriptome set (RPM >0). C, The proportion of genes expressed at very low (<1RPM), low (1-5RPM), moderate (5-20RPM) or high (>20RPM) levels in RNAseq sets. Abbreviations: MMC, megaspore mother cell; fTET, female tetrads; FM, functional megaspore selection stage (not collected); ES2n, mitotic embryo sac with 2 nuclei; ES4n, embryo sac with 4 nuclei; MES, mature embryo sac at anthesis, where antipodals have degenerated, containing the egg cell (EC), flanking synergids (Sy, not collected), and central cell (CC); Em, embryo; En, endosperm (not collected); PMC.E, early pollen mother cell; PMC.L, late pollen mother cell; mTET, male tetrads; MIC, uninucleate microspore; Bi, bicellular pollen stage; Tri, tricellular pollen stage, two sperm cells (SC) in the emerging pollen tube; MP-SC, mature pollen-sperm cell.

The events of fertilization initiate with pollen grain germination and emergence of the pollen tube on the stigma. We have observed that mitosis of the generative cell to form the two sperm cells predominantly occurs after pollen tube germination (see Results). Pollen tube growth through maternal tissues into the ovule, synergid puncture and pollen tube burst releases both sperm cells into the embryo sac. One sperm cell fuses with the egg to initiate embryogenesis and the other fuses with with the polar nucleus of the central cell to produce the triploid endosperm. Endosperm tissue is transient in copwpea as it is utilized during embryo development (Salinas-Gamboa et al., 2016); Fig. 1A).

Studies in *Arabidopsis*, rice, maize, *Boechera* and *Hieracium* have used LCM to isolate cells involved in sexual and asexual (apomictic) reproductive events to examine cell-type specific gene expression (Anderson et al., 2013, Kubo et al., 2013, Zhan et al., 2015, Belmonte et al., 2013, Okada et al., 2013). These analyses have typically focused on the expression in cells involved in specific aspects of the reproductive pathway due to the biological questions being addressed, and potentially, the laborious nature of the method. To our knowledge, a transcriptome series spanning reproductive events in isolated cells from the initiation of meiosis to seed initiation has not been generated in a species. This would enable direct comparisons of gene expression during male and female gametogenesis and seed development within the plant and also with pathways known to functionally regulate cell specification and reproductive development in other plants.

Here, we used LCM and additional tissue isolation procedures, to develop a suite of 13 cowpea reproductive cell-type transcriptomes spanning the temporal events of male and female gametophyte development, and early seed initiation in cowpea. *In vitro* and *in planta* analyses were used to verify transcriptome integrity. Following an analysis of global gene expression profiles across the transcriptome set, an *in silico* analysis of the expression of genes involved in epigenetic regulation, hormone biosynthesis and signal transduction, and cell cycle progression was undertaken as these pathways regulate reproductive aspects in other angiosperms. These analyses revealed commonalities between cowpea and other species with respect to some of these pathways, in addition to novel cowpea-related differences identifying genes and pathways as candidates for future functional testing.

## Results

### Cowpea IT86D-1010 leaf and reproductive cell-type transcriptomes and PCR verifications

A total of 13 reproductive transcriptomes were generated, in duplicate, using RNA extracted from individual reproductive cells that span male and female gametogenesis, and early seed initiation. Laser capture microdissection was used to generate 12 reproductive cell-type transcriptomes, while pollen and sperm cells were sampled separately (see below, Fig. 1A). Female gametophyte-related cell-type transcriptomes were generated for the megaspore mother cell (MMC) during meiosis I, the tetrad of post-meiotic megaspores (fTET), developing mitotic embryo sacs with two or four nuclei (ES2n, ES4n), and the mature embryo sac (MES) containing the egg cell (EC), the flanking synergids (Sy), and the central cell (CC), at the time of stamen emergence (anthesis). Individual EC and CC transcriptomes were also made. The early embryo (Em) transcriptome comprised tissue from developing zygotes to early globular stages of cowpea embryogenesis (Fig 1A).

Male gametophyte-related transcriptomes were generated for pollen mother cells at early and later stages of meiosis I (PMC.E, PMC.L), microspore tetrads (mTET), and uninucleate microspores (MIC). A pure sperm cell transcriptome was not generated. Analyses of cleared anthers indicated that 9.7% of mature pollen grains contained two sperm cells (n=500), indicating the final generative cell mitosis giving rise to the sperm cells primarily occurs following pollen tube germination. This is also observed in soybean and 70% of flowering plant species (Brewbaker, 1967, Williams et al., 2014, Haerizadeh et al., 2009, Wojciechowski et al., 2004). Anthers and pistils harvested at anthesis were processed to create a transcriptome referred to as mature pollen-sperm cell (MP-SC; see Methods for details). An additional transcriptome generated from young expanding cowpea leaves served as a control comparison for gene expression in cells not undergoing reproductive events.

A total of 74,839 genes were predicted in the IT86D-1010 genome (Spriggs et al., 2018) using Augustus gene prediction software when *Arabidopsis* genes were used as the training data set (Supplemental Data 1). Transcriptome sequences were aligned with the predicted genes in the IT86D-1010 cowpea genome to identify common and uniquely expressed genes across all datasets (Supplemental Table 1, Supplemental Data 2). The total number of uniquely expressed genes within each transcriptome ranged from 22,561 in the female megaspore tetrad (fTET) to 60,359 in the MP-SC sample (Fig. 1B). Only 3,769 of the genes expressed in the MP-SC transcriptome were not detected in at least one other cell-type. As a likely consequence of its isolation method, the high transcript diversity in the MP-SC transcriptome contrasts with that found in pollen and sperm cell transcriptomes of *Arabidopsis*, rice and maize, where the number of genes detected is reduced compared to sporophytic cell-types (Rutley and Twell, 2015).

Gene expression across all of the transcriptomes was normalized using reads per million (RPM) given the absence of a correlation between gene length and read alignment number. Investigation of unique gene expression levels within each transcriptome indicated that the majority of unique genes (68-85%) were expressed at very low to moderate levels (<1 to 5-20 RPM; Fig 1C). Principal component analysis (PCA) using the individual transcriptome replicates showed that most replicates clustered together. The transcriptomes from male and female reproductive cell-types co-segregated forming two distinct male and female clusters. The embryo transcriptomes associated with the female reproductive transcriptome cluster, while the leaf and MP-SC transcriptomes were outliers (Supplemental Fig.2).

**Figure 2.**
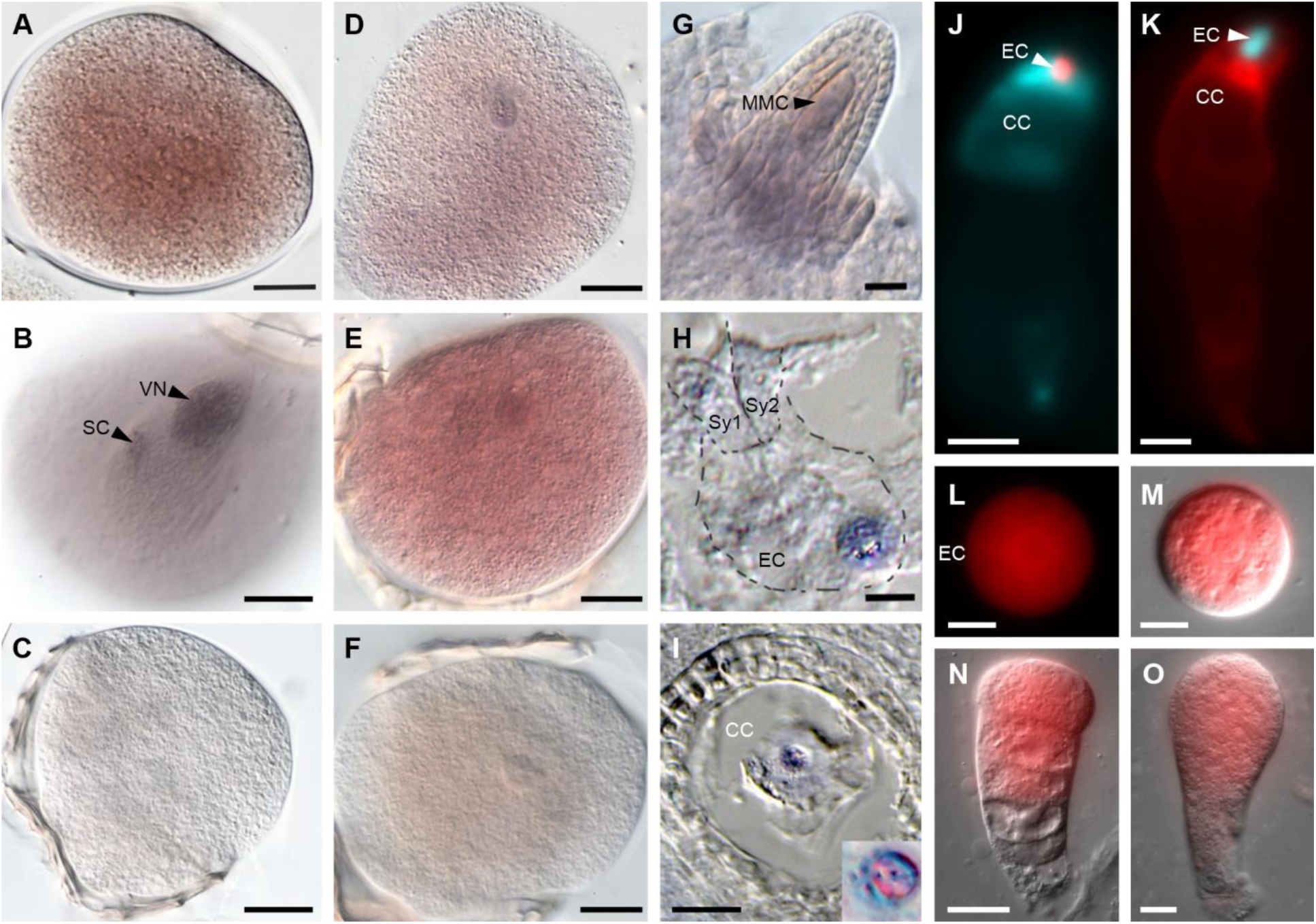
Gene expression in cowpea reproductive cell-types. A – J, Whole-mount *in situ* hybridization (WISH) in pollen grains of cowpea using four different antisense probes. A-C, *VuXTH6* mRNA in sperm cells A and vegetative nucleus B, with no signal in the sense control C. D and E, *VuXTH32* mRNA was detected in sperm cell and vegetative cell cytoplasm. F, *VuXTH32* sense controls. G, Whole-mount ovule undergoing megasporogenesis showing *VuKNU1* mRNA localized in the MMC, the nucellus and the inner integument. H, Paraffin section through a differentiated female gametophyte showing *VuRKD1* mRNA localization in the egg cell and synergids. I, Paraffin section through a differentiated female gametophyte showing *VuNB-ARC* mRNA localization in the central cell. Inset shows fused polar nuclei showing active transcription of *VuNB-ARC*. J, *AtRKD2_pro_*:*dsRED-Express* (red) in EC and *AtDD9_pro_:AmCyan1* (cyan) in the CC one day before anthesis. K, *AtEC1.1_pro_:AmCyan1* (cyan) in EC and *AtDD25_pro_:dsRED-Express* (red) in CC. L-M, *AtDD45_pro_*:*dsRED-Express* (red) in an isolated egg cell, N, two-celled proembryo and O, early embryo. Abbreviations: EC, egg cell; CC, central cell; Sy, synergids; SC, sperm cells; VN, vegetative nucleus, MMC, megaspore mother cell. Scale bars: A-G, L-M = 10 µm, H = 8 µm, I = 14 µm, J-K = 50 µm, N-O = 20 µm.

The quality of the generated transcriptomes was evaluated by examining if cell-type specific genes could be identified via *in silico* methods in the transcriptomes, and then by verifying gene expression by qRT-PCR, *in situ* and transgenic approaches. Candidate cell-type specific genes were initially identified using rsgcc software in the MMC, PMC, EN2n, En4n, MES, CC, ES, Em and sperm transcriptomes (Fig. 1A). Candidate genes identified ranged in number from 126 in early pollen mother cells to 25,986 in the MP-SC transcriptome (Supplemental Data 3 and 4). A filtering approach based on identification of genes expressed at least five times higher in the CC, EC or MP-SC transcriptomes relative to all others resulted in 10, 34 and 31 candidate specific genes, respectively (Supplemental Table 2).

Two of the cowpea egg cell-specific candidates identified using the filtering approach were homologues of the *Arabidopsis* egg cell-specific genes *AtEC1.1*, (Sprunck et al., 2012) and *AtRKD1* (Koszegi et al., 2011) (*VuEC1.1_1* and *VuRKD1*; Table 1). Therefore, cowpea homologues of 11 additional *Arabidopsis* reproductive cell-type expressed genes were identified for verification analyses. They included *VuKNU-like1* and *2*, putative cowpea homologs of the *Arabidopsis KNUCKLES* gene expressed during male and female gametogenesis (Payne et al., 2004, Tucker et al., 2012) and genes expressed in specific male and female gametophytic cell-types (Table 1; Supplemental Table 3). Twenty of the 28 (71%) candidate cell-type specific genes tested by qRT-PCR, were confirmed as significantly enriched using cDNA isolated from tissues at comparable developmental stages from which the LCM samples were obtained in addition to other tissue types (Table 1, Supplemental Table 4).

**Table 1.**
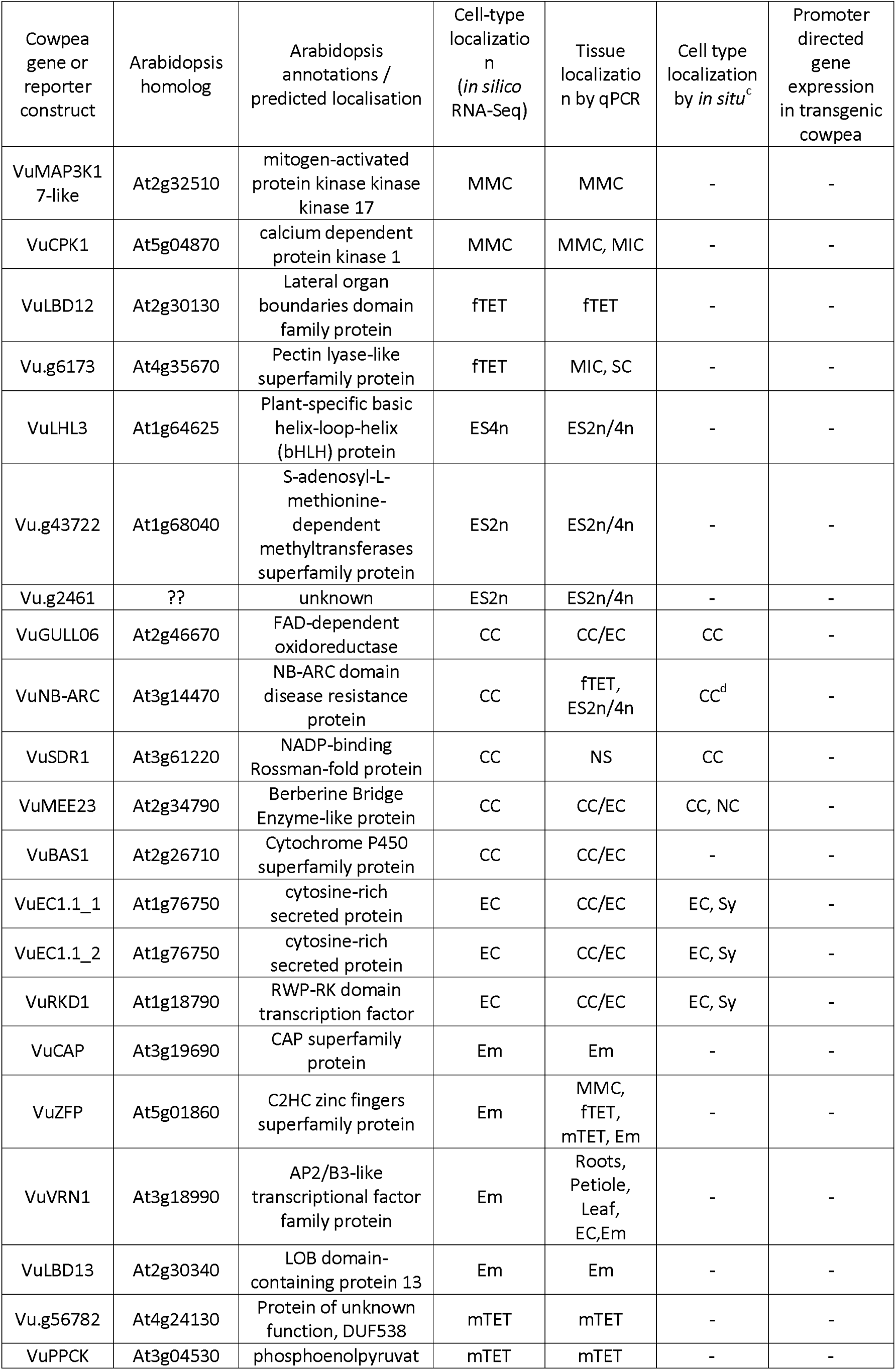
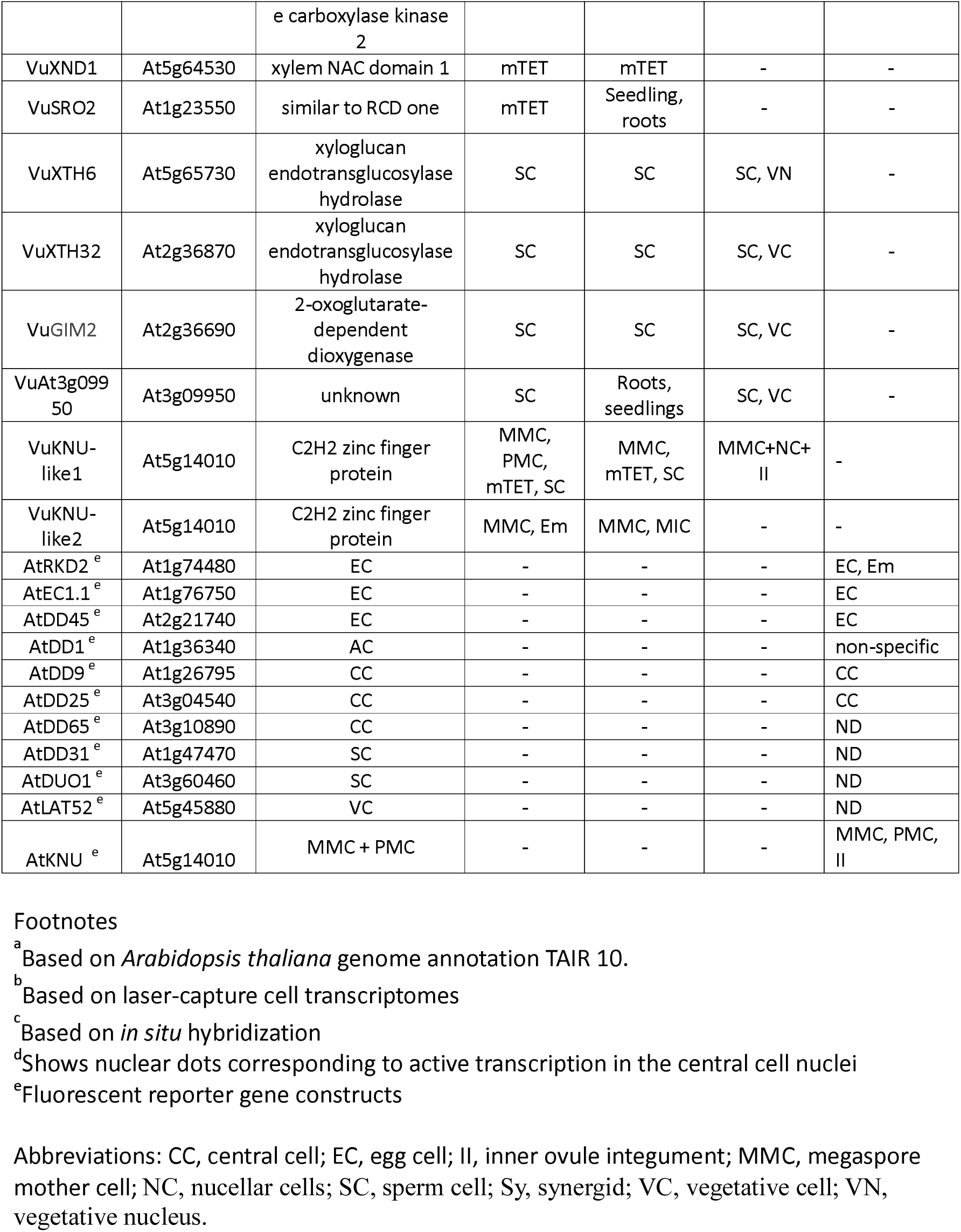
Genes and promoters tested for cell-type specificity by qPCR, *in situ* hybridization and transformation in cowpea.

**Table 2.** The number of genes upregulated in each cell-type compared to the average expression in all cell-types.

| Cell type | Number of genes differentially upregulated |
| --- | --- |
| MMC | 5878 |
| fTET | 2316 |
| ES2n | 2627 |
| ES4n | 1423 |
| CC | 1897 |
| EC | 1550 |
| MES | 3376 |
| Em | 5051 |
| PMCE | 1282 |
| PMC L | 2653 |
| mTET | 3743 |
| MIC | 3101 |
| sperm | 38691 |
| Leaf | 9140 |

### Verification of cell-type specific expression using in situ hybridization and plant transformation

The expression of 11 candidate cell-type specific genes verified using qRT-PCR analyses above was examined by *in situ* hybridization (ISH), using gene-specific digoxygenin-labelled probes (Fig. 2 and Supplemental Fig. 4). ISH was performed on mature pollen grains to examine expression of four candidate sperm-cell specific genes, as we had previously determined that approximately 10 percent of mature pollen grains contain two sperm cells (Table 1). *VuXTH6* mRNA was detected at high levels in the sperm cells and in the vegetative nucleus, with low levels detected in the vegetative cell cytoplasm (Fig. 2A-C). By contrast, mRNA for the other three examined genes was detected in the sperm cells, the vegetative nucleus and at high levels in the vegetative cell cytoplasm (Fig. 2D-F and Supplemental Fig. 4A-F).

**Figure 3.**
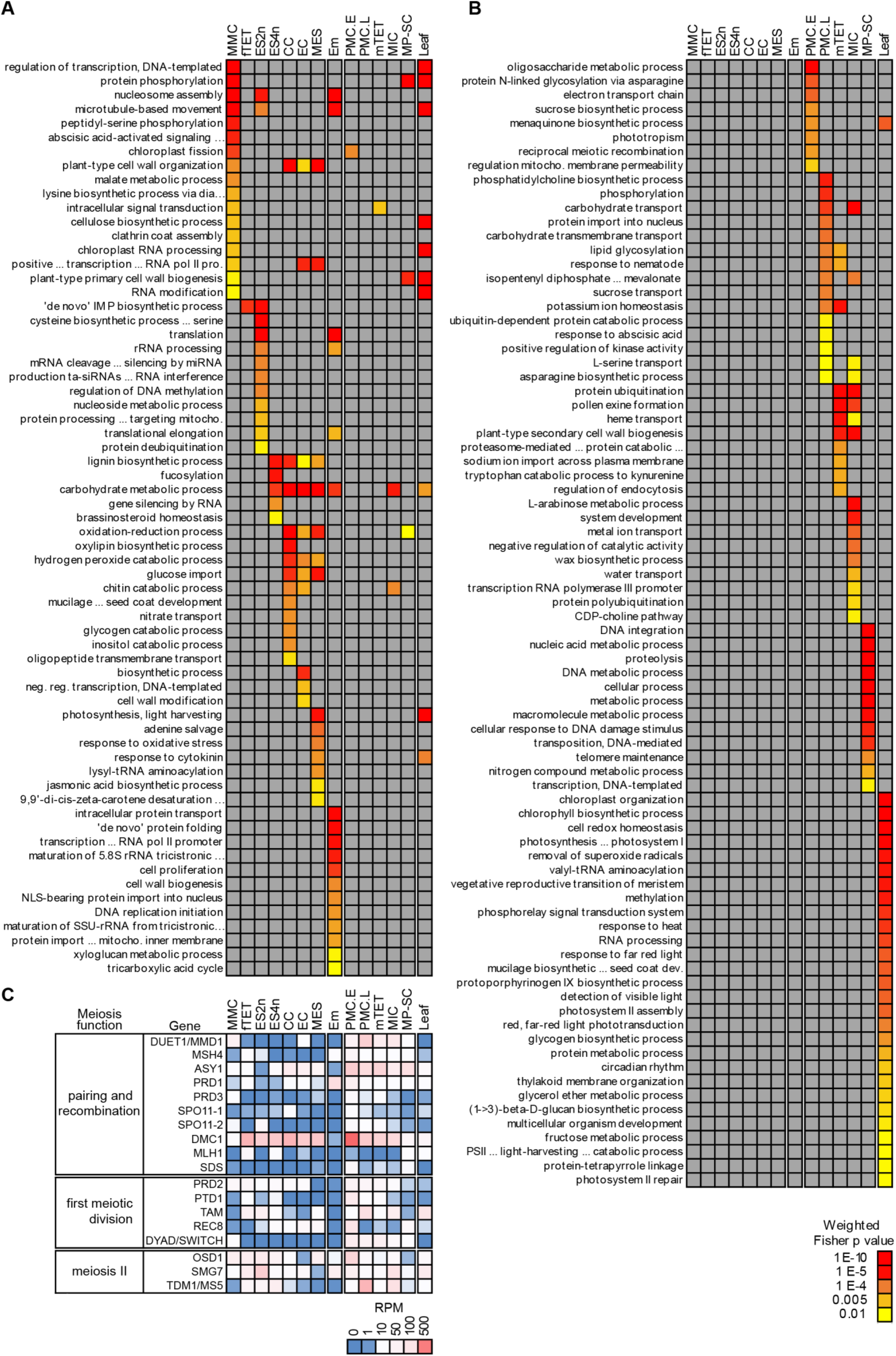
Differentially enriched Biological Process GO terms and expression of meiosis genes during gametogenesis. A, Enriched GO terms primarily in female gametophyte and early embryo transcriptomes. B, Enriched GO terms primarily in the male gametophyte and leaf transcriptomes. The statistical significance of the GO enrichment is represented according to the scale of weighted Fisher p value shown at the bottom right. C, Expression of genes involved in meiosis in the generated cowpea transcriptomes (See also Supplemental Tables 7 and 8). Expression levels relate to the RPM scale.

**Figure 4.**
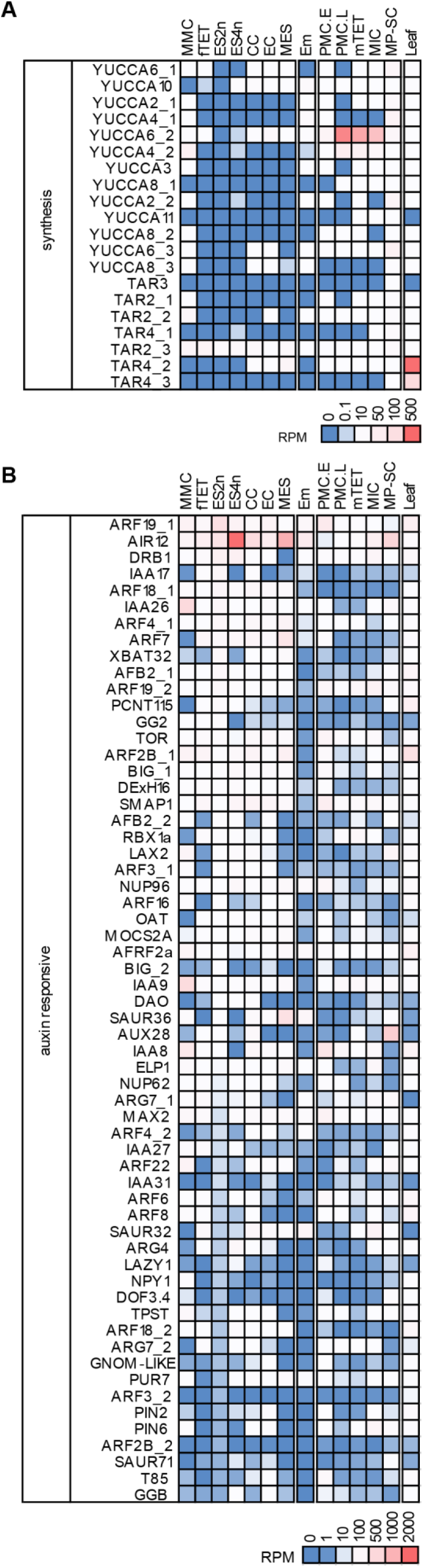
Expression of auxin biosynthesis and auxin responsive genes during meiosis and gametophyte development. A, Expression of genes belonging to the YUCCA and TAR auxin biosynthesis families. B, Expression of 60 auxin responsive genes sorted in decreasing order of expression level in the 2-nucleate embryo sac. Expression levels are related to the RPM scale with each respective panel. (See also Supplemental Table 9).

A combination of whole-mount and section-based *in situ* hybridization was used in developing cowpea ovules used to examine expression of seven genes during female gametogenesis. Transcripts of *VuKNU-like1*, the candidate cowpea homolog of the *Arabidopsis KNUCKLES* gene were enriched in the megaspore mother cell; however, expression was also observed in nucellar and inner integumentary ovule cells (Fig 2G). Transcripts from the candidate egg cell specific *VuEC1.1_1* and *VuRKD1* genes (Table 1) were highly enriched in the egg cell, in comparison to the rest of the embryo sac and ovule. In both cases, mRNA localization was also evident in the synergids, but not in the central cell (Fig. 2H; Supplemental Fig. 4G-J). Three of the four candidate central cell genes tested by *in situ* hybridization, displayed specific mRNA localization in the central cell (*VuSDR1, VuGULL06*; Table 1; Supplemental Fig. 4N and O, Fig 2I). *VuNB-ARC*, mRNA was evident in a nuclear dot patterning in the cowpea central cell nucleus (Fig. 2I). Nuclear dots are tightly associated with nascent transcripts of actively expressed genes in the central cell of *Arabidopsis* (Vielle-Calzada et al., 1999), and in embryonic cells of *Drosophila melanogaster* (Shermoen and O’Farrell, 1991). *VuMEE23* mRNA was localized in the central cell and in adjacent ovule cells (Table 1).

As eleven of the cowpea genes predicted to be expressed in specific reproductive cell-types showed sequence similarity to previously described *Arabidopsis* cell-type expressed genes, cowpea plants were transformed with constructs comprising *Arabidopsis* (*At*) promoters of these genes fused to fluorescent reporters to examine conservation of targeting to specific reproductive cell-types (Table 1; Supplemental Table 3). Six of the tested promoters directed similar spatial and temporal patterns in cowpea as observed in *Arabidopsis*. The remaining five showed either non-specific or no detectable expression in transgenic cowpea (Table 1; Fig. 2 and Supplemental Fig. 4). The *AtKNU* promoter directed expression to the MMC and PMC cells, as published previously (Tucker et al., 2012) but expression was also evident in the chalazal region of the cowpea ovule integument (Supplemental Fig. 5F). *In situ* hybridization using probes to detect native cowpea *VuKNU* expression confirmed expression in the ovule integument (Supplemental Fig.4K). The *AtDD45* promoter directed expression to the egg cell (Fig. 2J) and both *AtDD9* and *DD25* promoters enabled central cell expression (Fig. 2K). The *AtEC1.1* and *RKD2* promoters directed reporter expression to egg cells of transgenic cowpea (Fig. 2K-M). Expression of fluorescent reporters from *AtRKD2* and *DD45* promoters was also observed in early cowpea embryos (Fig. 2N-O).

**Figure 5.**
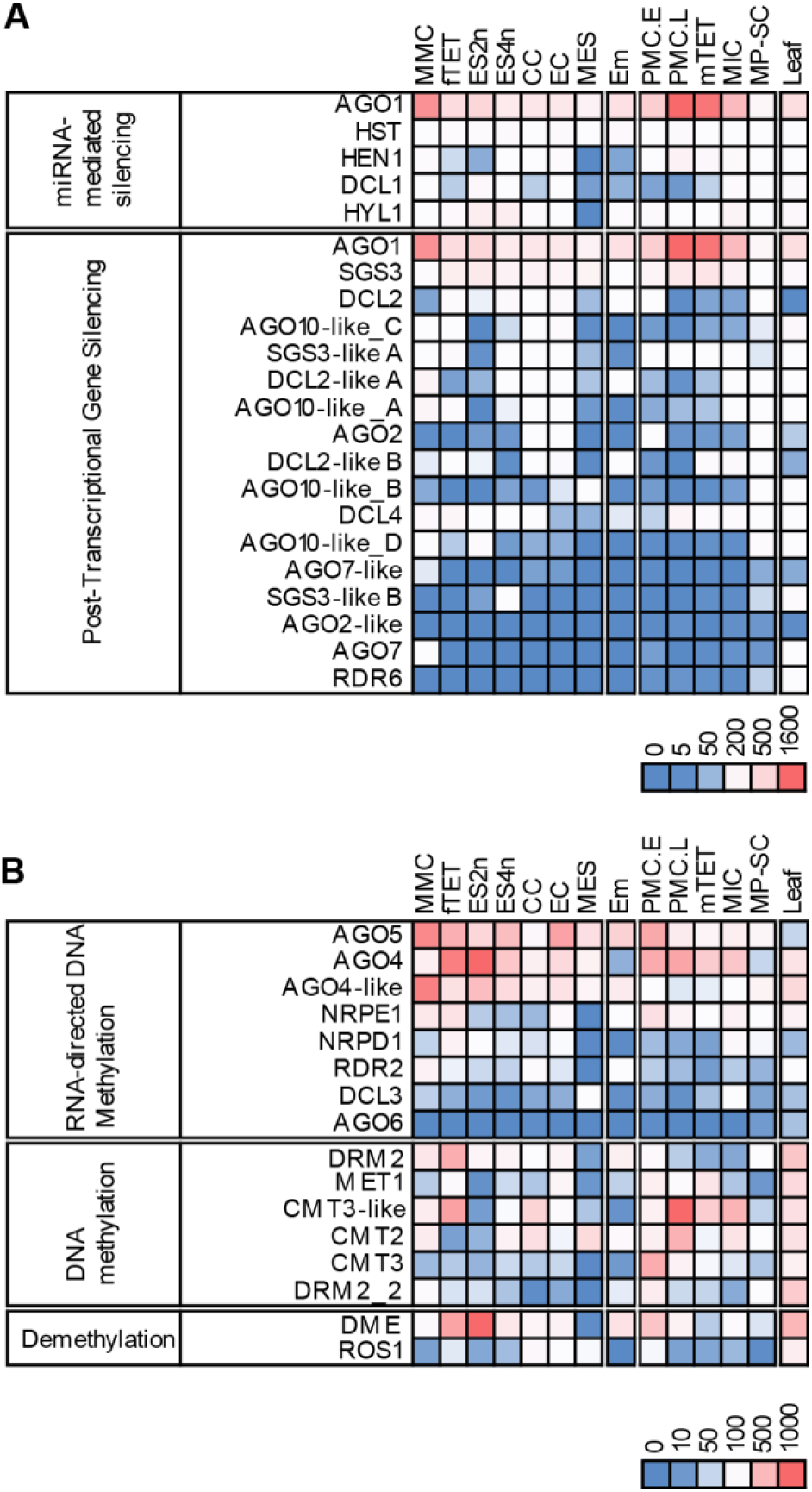
Heatmaps representing the expression of selected genes involved in RNA-mediated gene silencing and DNA methylation. A, Expression of genes involved in the miRNA-mediated gene silencing pathway and in PTGS. B, Expression of genes that interact with RNA and are involved in RdDM, DNA methyltransferases that mediate RdDM and DNA glycosylases that mediate demethylation of DNA. Expression level is according to the RPM scales for each panel. (See also Supplemental Tables 12 and 13).

Given the low efficiency and lengthy nature of the cowpea transformation protocol used, we also examined if upstream promoter regions of 11 of the cowpea candidate cell-type specific genes identified from *in silico* analyses would direct the same cell-type expression in *Arabidops*is. Genomic fragments of 0.8 to 3.2 kb, corresponding to their upstream putative promoter regions were fused to the *uidA* (GUS) reporter gene and transformed into the *Arabidopsis* Columbia (*Col-*0) ecotype. For each transcriptional fusion, at least 15 independent transformants were recovered and histologically analyzed to examine gene expression patterns during male and female gametogenesis. GUS activity was not detected in reproductive organs of *Arabidopsis* in seven of the 11 tested constructs. However, a 1,429 bp region upstream of *VuKNU-L1* drove GUS expression in the *Arabidopsis* MMC and adjacent sporophytic ovule cells, in a pattern similar to the *VuKNU-L1* mRNA ISH localization pattern in cowpea (Supplemental Fig. 6A and Supplemental Table 21). The promoter fragments from the *pVuXTH32* and *pVuGIM2* genes drove spatial expression in both pollen sperm and vegetative cells as observed in cowpea ISH experiments (Supplemental Fig. 6F-G). By contrast, the promoter of *pVuRKD1* drove expression in the egg cell, and in both sperm and vegetative cells in 5 of the 15 *Arabidopsis* transformants analyzed (Supplemental Fig. 6B-D). In seven of the transformants, expression was seen only in sperm cells (Supplemental Fig. 6C). Finally, the promoter of the cowpea gene *VuGULL06*, a candidate selected for potential central cell expression drove expression in the *Arabidopsis* egg cell and synergids (Supplemental Fig. 6E), suggesting that in some cases cowpea promoters can be influenced by heterologous factors intrinsic to the *Arabidopsis* genome.

**Figure 6.**
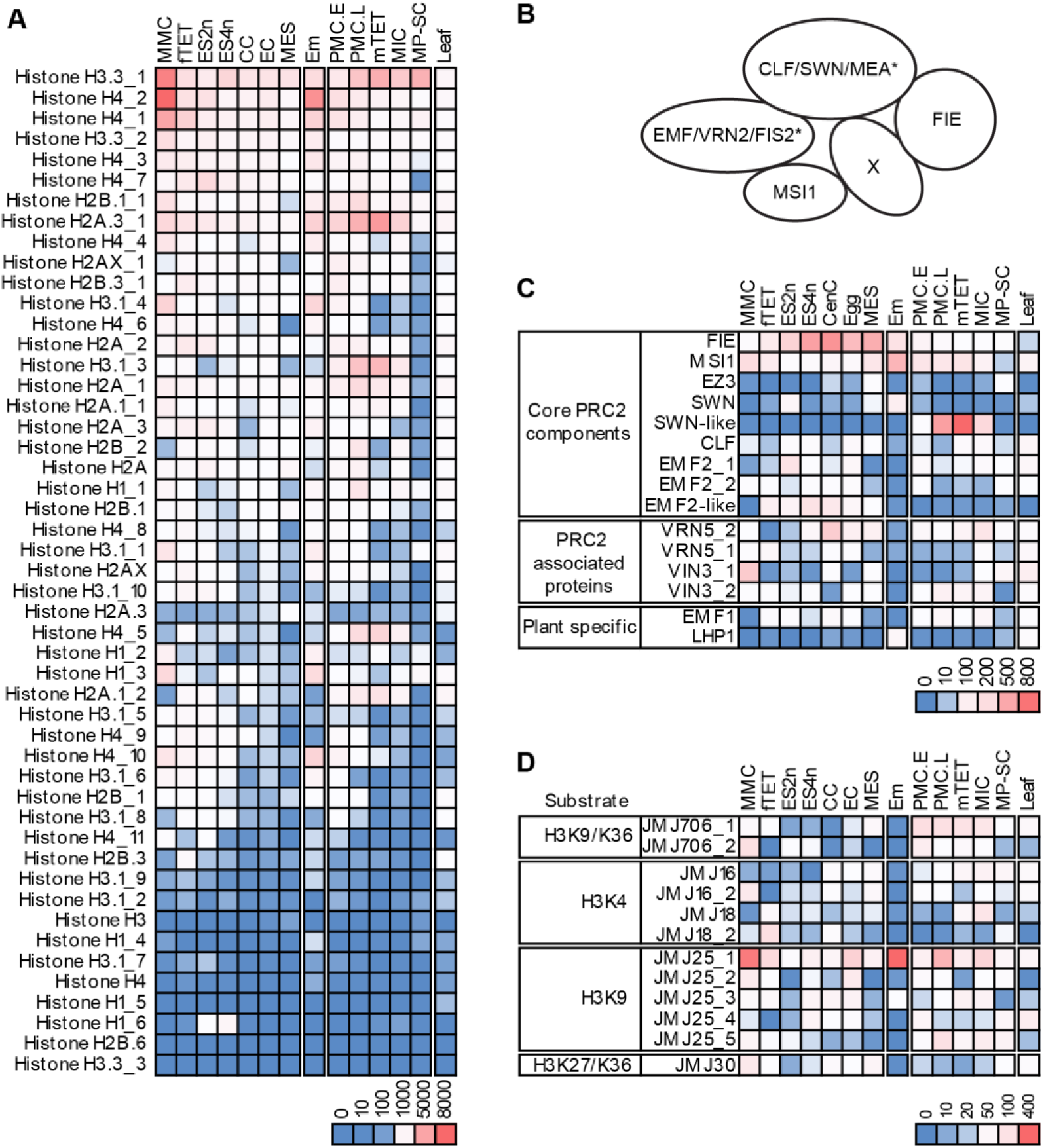
Histone gene expression and the expression of selected genes involved in histone modification. A, Expression of genes belonging to the core histone family (See Supplementary Table 18). B, Expression of selected PcG genes involved in histone methylation resulting in gene silencing (See Supplementary Table 19). C, Known protein interactions forming Polycomb-group repressive 2 complex types. Accessory proteins (X) are often associated with PRC2 complex function. Asterisks indicate *Arabidopsis* specific proteins. MEA and *Arabidopsis* counterparts, SWN and CLF are SET domain proteins and homologs of *Drosophilia* Enhancer of Zeste. FIS2 and others listed are VEFS-box proteins. FIE and MSI1 are WD40 repeat protein containing proteins (Mozgova and Henning, 2015). D, Expression of selected JMJ family histone demethylase genes and demethylation targets, at left (See also Supplemental Tables 14, 15, 16). Expression levels relate to the RPM scales associated with each panel.

The promoter analyses indicated that there is some conservation in regulators enabling reproductive cell-type specific expression in cowpea and Arabidopsis given the conservation of gene expression directed to common cell types in transgenic studies, in some cases. Taken together, these verification analyses collectively indicted that the developed transcriptomes were a reliable resource for further *in silico* interrogation of gene expression profiles defining specific reproductive cell-types during male and female gametogenesis and early cowpea embryogenesis.

### Enriched biological processes in cowpea reproductive transcriptomes

Differentially expressed (DE) genes were identified in each transcriptome type to examine gene expression changes occurring during the temporal events of male and female gametogenesis and early embryogenesis (Supplemental Data 4). Percentages of differentially expressed genes varied widely from 3.4% to 25% amongst the transcriptomes (Supplemental Table 5). Comparisons between the cowpea female reproductive cell-type transcriptomes generated using LCM and those previously generated using LCM in *Arabidopsis* revealed low levels of common genes. Comparisons were made between the cowpea data set and those generated in *Arabidopsis* for the MMC (Schmidt et al., 2011), ovule nucellus, early mitotic female gametophytes (Tucker et al., 2012) and the egg and central cell (Wuest et al., 2010, Steffen et al., 2007). The highest similarity, with a 24% overlap, was observed between the cowpea MMC dataset and the *Arabidopsis* female nucellus plus MMC data set (Tucker et al., 2012); Supplemental Table 6). The low levels of common genes observed may reflect differences in species-specific genes, staging, and methods used to generate transcriptomes and identify expressed genes.

To obtain an overview of the major biological processes occurring during gametogenesis and seed initiation, each transcriptome was initially surveyed by examining significantly enriched gene ontology (GO) terms in the biological processes’ category (Fig. 3A-B). Examination of the control expanding leaf transcriptome showed that it was, unsurprisingly, enriched in GO terms related to light transduction, chlorophyll, photosynthesis, and sugar metabolism (Fig. 3). Each cowpea reproductive transcriptome was characterized by a relatively distinct set of significantly enriched biological process terms, supporting the cytologically distinct nature of the cell-types collected (Fig. 3A-B).

Meiosis and recombination, and cell wall biogenesis terms, which characterize the functional and cytological events of male and female sporogenesis in angiosperms, were found in cowpea meiotic cell transcriptomes. Terms related to a range of pathways involved in epigenetic regulation were also prevalent in transcriptomes related to the elaboration of the female and male gametophytes (Fig. 3A-B). Regulation by small RNAs and global changes in DNA and histone methylation modifications influence determination, selection and development of gametophytic cells (Tucker et al., 2012, Olmedo-Monfil et al., 2010). They prevent transposon activation during male and female gametogenesis (Slotkin et al., 2009, Schoft et al., 2011, Pillot et al., 2010, Ibarra et al., 2012, Martinez et al., 2016). In *Arabidopsis*, Histone H3 methylation (at lysine (K) 27) by the *Polycomb*-Group protein repressive complex 2 (PRC2), results in gene silencing and regulates *Arabidopsis* male gametophyte formation and female gametophyte arrest, pre-fertilization (Mozgova and Hennig, 2015, Grossniklaus and Paro, 2014, Wang and Kohler, 2017, Kawashima and Berger, 2014).

The plant hormone auxin is involved in a range of reproductive events in angiosperms, including initiation of seed formation and embryo patterning. Auxin-related terms were very evident in this global analysis. Interestingly, the GO enrichment analysis suggested potential involvement of other plant hormone pathways at particular cowpea developmental stages. For example, the abscisic acid signalling pathway featured in the female MMC and male late PMC transcriptomes. Brassinosteroid-related terms were evident in dividing embryo sacs with 2 and 4 nuclei, and terms related to jasmonic acid biosynthetic processes and responses to cytokinin were evident in the mature cowpea embryo sac (Fig. 3A and Supplemental Data 6).

In order to gain greater understanding of reproduction-related gene expression pathways that cowpea may, or may not have in common with other angiosperms, we profiled the transcriptome set for the expression of cowpea homologues of *Arabidopsis* genes involved in meiosis, auxin biosynthesis and signal transduction, epigenetic pathways and cell-cycle progression. The results of these analyses, in the sections below, collate examined gene expression profiles during male and female gametogenesis, pre-fertilization female gametophyte arrest and early embryogenesis.

### Gene expression during temporal events of cowpea male gametophyte formation

The expression of 18 cowpea meiosis-related gene homologues during male gametogenesis is shown in Fig. 3C. The profiled genes have putative functions in chromosome pairing, homologous recombination, and subsequent meiotic events (see Supplemental Table 7 for functional information and references). A number of these genes were clearly expressed in non-meiotic cell-type transcriptomes and in leaf (Fig. 3C), consistent with expression data from other species (Klepikova et al., 2016).

Analyses of the expression of 276 auxin biosynthesis and signal transduction pathway genes in male gametogenesis showed high levels of the cowpea *YUCCA6-*like auxin biosynthesis gene in transcriptomes of cowpea late pollen mother cells, tetrads and microspores (Fig. 4A, Supplemental Table 9). This reflects the requirement of *YUCCA6* and auxin production in corresponding cells for viable *Arabidopsis* pollen development (Yao et al., 2018). Cowpea homologs of the *Arabidopsis ARF19*, *AIR12* and *DRB1* auxin-responsive genes were also upregulated during male gametophyte formation (Fig 4B). In *Arabidopsis*, ARF19 enables auxin-activated transcriptional responses to phosphate starvation (Huang et al., 2018). AIR12 is thought to link auxin signal transduction and reactive oxygen signalling (Gibson and Todd, 2015) and DRB1 is involved in cleavage to produce miR399 phosphate homeostasis regulation (Pegler et al., 2019).

Cowpea *ARGONAUTE* (*AGO*) genes were identified because AGO proteins are central to the small RNA-mediated processes of Post-transcriptional Gene Silencing (PTGS), microRNA (miRNA) mediated gene silencing and RNA-directed DNA methylation (RdDM). The 13 cowpea *AGO*-like genes identified showed close phylogenetic relationships with known *AGOs* in other plants, although *AGO8* and *AGO9* were not evident in the cowpea genome (Supplemental Fig. 7).

**Figure 7.**
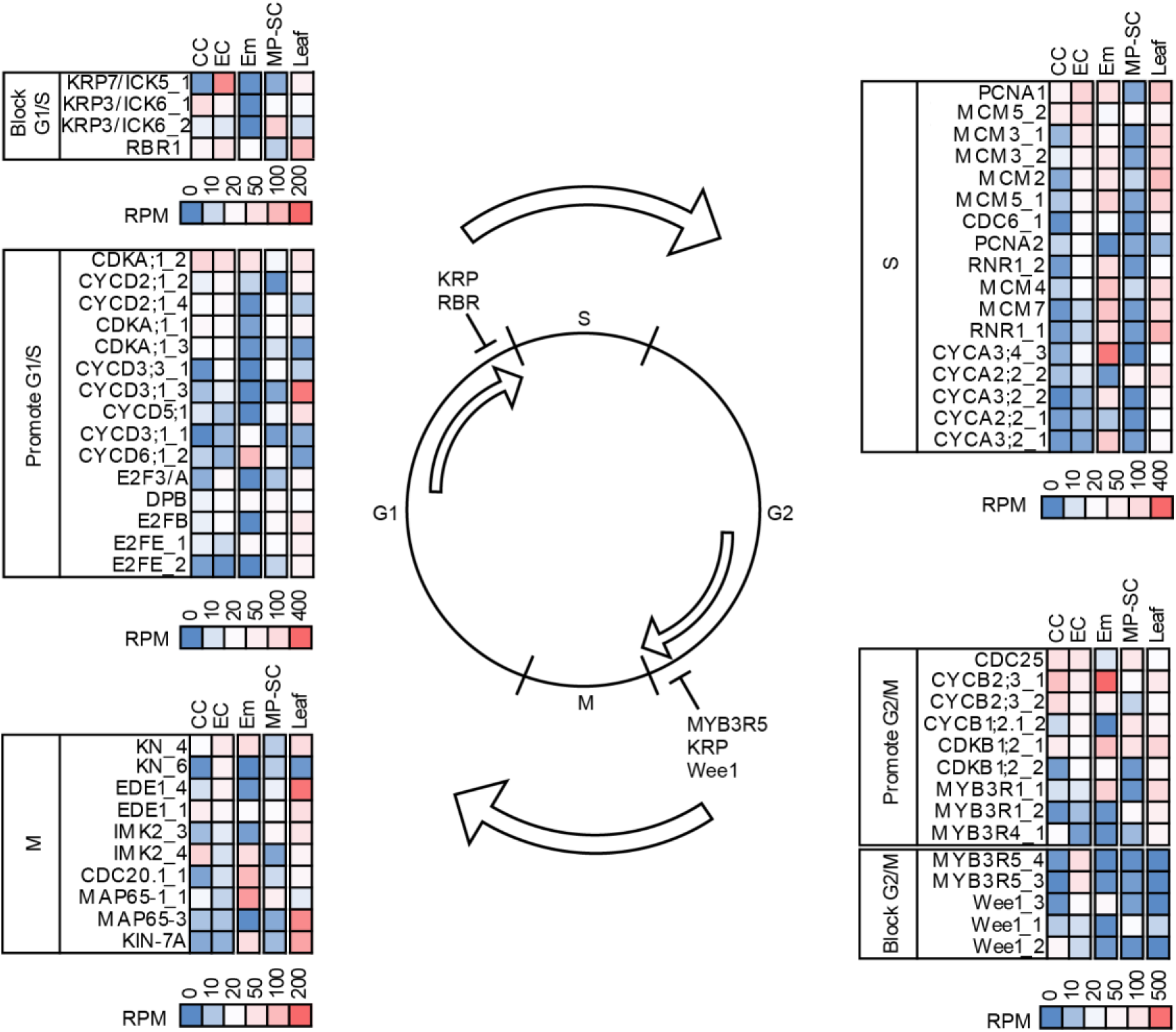
Expression of selected cell cycle genes in egg, central cell, mature pollen-sperm and leaf transcriptomes. Cell cycle genes are grouped according to their function relating to roles in promoting or blocking progression through various cell cycle phases. Relative expression level is indicated according to the RPM scale shown at the bottom of each panel. (See also Supplemental Tables 10 and 11).

Analyses of cowpea homologs of genes required for the 21 nt producing *Arabidopsis* microRNA pathway showed that *AGO1*, *HEN1*, *HYL1* and *HST1* were co-expressed during meiotic and mitotic stages of male gametophyte development. This suggested potential conservation of this pathway with a functional role during cowpea male gametophyte elaboration (Fig. 6A).

By contrast, the case for a functional *Arabidopsis*-like PTGS pathway operating during male gametogenesis appeared doubtful. The *Arabidopsis* PTGS pathway typically produces 21-24 nt small interfering (si) RNAs. It requires the function of AGO1, RDR6, SGS3 and DCL4 as core protein components. However, AGO2, AGO7, and DCL2 can act with partial redundancy with their respective paralogs (Borges and Martienssen, 2015). Homologs of all core *Arabidopsis* PTSG components including two *AGO2*, two *AGO7*, four *AGO10* and three *DCL2*-like genes were identified in the cowpea genome. All of the core components required for *Arabidopsis*-like PTGS were expressed during male gametogenesis except for *RDR6*, which was undetectable or expressed at very low levels in all male cell-transcriptome types and at low levels in the mature pollen-sperm cell sample. By contrast, moderate *RDR6* expression was evident together with similar or higher expression of the PTSG components in the leaf transcriptome (Fig. 6A and Supplemental Table 10 and 11). The typical *Arabidopsis*-like PTSG pathway may not be active during cowpea male gametogenesis.

The *Arabidopsis* RdDM pathway results in the production of 24nt siRNAs and site-specific *de novo* methylation at their target DNA sites by DNA methyltransferases MET1 and DRM2 (Borges and Martienssen, 2015). Genes required for RdDM including *AGO4*, *AGO5*, *DCL3*, *RDR2*, *NRPD1* and *NRPE1* (encoding the largest subunits of POL IV and V respectively) were detected at various stages during male gametogenesis. Transcript levels of *NRPD1*, *RDR2* and *DCL3* were relatively low. However, highest transcript levels of DNA methyltransferase *MET1*, *DRM2_1* and *DRM2_2* were evident during male meiosis, decreasing during the mitotic events of gametogenesis (Fig. 6B and Supplemental Table 11). This suggests an *Arabidopsis*-like RdDM pathway is likely to be functional during cowpea male gametogenesis.

Other DNA methyltransferases including CMT2 and CMT3 DNA can maintain CHG/CHH and CHG methylation, respectively, independently from the RdDM pathway in *Arabidopsis*. Cowpea homologs of both genes were strongly expressed during male meiosis and mitosis (Fig. 5B). Interestingly, high transcript levels from the DNA glycosylase *DME* that mediates DNA demethylation by removing methyl-cytosine and replacing it with unmethylated cytosine in *Arabidopsis* (Penterman et al., 2007) were found during early cowpea PMC meiosis (Fig. 6B and Supplemental Table 13). Collectively, these data suggest multiple pathways inducing dynamic DNA methylation changes may operate during cowpea male gametogenesis.

Changes in chromatin histone composition are known to occur during pollen development in *Arabidopsis*. A total of 44 cowpea histone genes belonging to the core histone families H2A, H2B, H3 and H4 were identified in cowpea (Talbert et al., 2012); Fig. 6A and Supplemental Table 12), and their expression profiled during male gametogenesis. A prevalence of H3.3 histone transcript variants are evident in mature *Arabidopsis* pollen and H3.1 variants were absent (Borg and Berger, 2015). Analysis of histone gene transcription during cowpea male gametogenesis showed high levels of the core histone H3.3 transcripts throughout male gametogenesis, including the mature pollen-sperm cell sample. H3.1_3 histone transcript levels were also detected during male gametogenesis but were undetectable in the mature pollen-sperm cell sample, as was a range of other cowpea histone transcripts (Fig. 6A). These data suggest conservation of transcriptional patterns of H3.3 and 3.1 expression during pollen maturation in cowpea and *Arabidopsis*.

Epigenetic processes involving the Polycomb-group repressive complex 2 (PRC2), which applies a repressive trimethylation mark on lysine 27 of histone H3 (H3K27me3) regulate a number of events in the plant lifecycle. Proteins forming the PRC2 complex vary in relation to these functional contexts (Fig. 6B). In *Arabidopsis*, the PRC2 VRN complex, containing CLF/SWN, VRN2 and FIE proteins, functions during vegetative development and the floral transition, whereas the EMF complex, containing FIE, MSI1, CLF/SWN and EMF2, functions during inflorescence development and flower organogenesis. The PRC2 FIS complex, containing MEA/SWN, FIS2, FIE and MSI1, functions to silence target genes during early seed development and gametogenesis (Pien and Grossniklaus, 2007). Cowpea homologs of *Arabidopsis FIE, MSI1, EMF2, VRN5, VIN3* and *EMF1* genes were co-expressed during male gametogenesis. (Fig. 6C). High levels of a cowpea *SWINGER*-like (*SWN*-like) transcript were observed during late pollen mother cell meiosis and microspore formation. The expression patterns of the PRC2 complex-like genes during male gametogenesis in cowpea suggest there is a potential for the formation and function of novel, possibly multiple PRC2 complexes resembling modified versions of the EMF and VRN PRC2 types during male gametogenesis.

Jumonji-domain proteins (JMJ) function to demethylate histones, antagonizing the establishment of histone methylation. *JMJ706_1*, *JMJ16* and *JMJ25_1* and *JMJ25_5*-like genes, potentially encoding proteins which demethylate histone methylation marks, other than H3K27 (Fig. 6D) were expressed at elevated levels during cowpea male gametogenesis (Fig. 6D). In rice, JMJ706 regulates transcription factor expression required for spikelet development, floral morphology and organ number (Sun and Zhou, 2008) *Arabidopsis* JMJ16 represses leaf senescence (Liu et al., 2019) and JMJ25 functions in poplar to remove H3K9me2 marks and modulate anthocyanin biosynthesis (Fan et al., 2018).

### Gene expression during temporal events of cowpea female gametophyte formation

Meiosis-related genes were expressed at a number of stages during female gametophyte development as also observed during male gametogenesis. Transcript levels of meiosis genes were evident at lower levels throughout female gametogenesis (Fig. 3C). Auxin biosynthesis gene transcripts were detected from *YUCCA* and *TAR*-like genes during cowpea megaspore mother cell meiosis (Fig. 4A). In contrast with male gametophyte development, auxin biosynthesis gene transcripts were generally undetectable in developing cowpea female gametophytes in most of the post-meiotic developmental stages. Exceptions were a *TAR2_3*-like gene expressed at relatively low levels throughout the stages of female gametogenesis, which was also expressed throughout male gametogenesis, during early embryogenesis and in leaf (Fig. 4A). *YUCCA6, 8, 10* and *TAR4_2*-like genes were co-expressed in the EC and CC with *TAR2_2* showing higher relative EC expression (Fig. 4A; Supplemental Table 9).

In direct contrast to the subdued nature of auxin biosynthesis gene expression in developing female gametophytes, a total of 47 genes annotated generally as “auxin-responsive” were expressed with RPM counts over 10 in cell-types of the developing female gametophyte (Fig. 4B and Supplemental Table 9). These genes included 15 *AUXIN RESPONSE FACTORS* (*ARF*s and five *AUX-IAA* genes, five auxin transporters belong to the *LAX* and *BIG* gene families, two polar auxin transport *PIN* homologs and three *SMALL AUXIN UP RNA* (SAUR) which are induced in response to auxin (Fig. 4B and Supplemental Table 9). The auxin induced *AIR12*-like gene was expressed at high levels in both the 4 nucleate and the mature embryo sacs, which suggests that regulation of reactive oxygen species may be important if the function of *Arabidopsis* AIR12 is conserved in cowpea.

The expression of gene components required for *Arabidopsis*-like micro-RNA-mediated silencing was evident during female gametophyte development. Low levels of *DCL1*, *HYL* and *HEN* were observed in the mature embryo sac. The relative expression of *DCL1* was approximately five times higher in the EC than CC (Figure 5A and Supplemental Table 11). These observations suggest that miRNA induced gene silencing is ongoing during the events of meiosis and mitosis during female gametogenesis. Potentially, miRNA production may be more active in the egg than the central cell in mature embryo sacs.

Examination of expression of homologs of genes involved in the *Arabidopsis* PTGS pathway showed non-detectable *RDR6* expression as also observed during in male gametogenesis. Thus either the typical PTGS pathway described for *Arabidopsis* is not active within male and female cowpea reproductive cell types examined, or other cowpea-specific elements substitute for generation of 21-24 nt siRNAs (Fig. 5A).

Expression of the RdDM pathway components during female gametogenesis suggested functional gene silencing is active during meiosis and mitotic events. High expression levels of *DME* were also observed in the post-meiotic megaspore tetrad and bi-nucleate mitotic embryo sacs, suggesting methylation turnover is required for the initiation of the mitotic embryo sac progression (Fig. 5B). Interestingly, expression of both *AGO4* and *AGO5* were approximately two times and five times higher in the EC than in the CC, respectively. In addition, *MET1* and *DRM2_1* were expressed at three-fold and 1.5-fold higher levels in the EC than CC, respectively. This suggests relatively lower gene silencing activity via the RdDM pathway in the CC than in the EC (Fig. 5B and Supplemental Table 11). Expression of homologs of *CMT2* and *CMT3-B* which maintain methylation independently from RdDM were approximately three times lower in egg than in central cell (Fig. 5B and Supplemental Table 11). These results suggest a hypothesis that possibly *de novo* DNA methylation via the RdDM pathway may be reduced in the cowpea CC compared to the EC, while maintenance of CHG and CHH methylation may be higher in the cowpea CC than the EC. DNA methylation turnover in the EC and CC may be mediated by *DME* and ROS as indicated by moderate levels of expression in these cell-types (Fig. 5B).

Most core histones including *H3.3-1* were expressed at moderate levels during female development in cowpea. (Fig. 6A and Supplemental Table 14). The MMC showed higher relative levels of histone *H3.3_1*, *H4_1, 2*, and *3.1_1* transcripts. *Histone3.3* variants were expressed throughout gametogenic development, as were a number of *H3.1* variants which contrasts with the trend observed in male gametogenesis (Fig.6A). High transcript levels of the JMJ25-like gene were evident at MMC meiosis, together with moderate levels of the JMJ706 H3K9 demethylase as seen in late pollen mother cell meiosis, suggesting a possible requirement for H3K9 demethylation in both male and female cowpea meiosis. Expression of the JMJ histone demethylase JMJ30, which removes H3K27me2/3 deposited by PRC2 complexes was higher in the female gametophyte than the male (except for the MP-SC sample; Fig. 6D; Supplemental Table 14).

In *Arabidopsis*, the PRC2 FIS complex, containing MEA/SWN, FIS2, FIE and MSI1, functions to silence target genes during embryo sac maturation and early seed development (Mozgova and Hennig, 2015, Grossniklaus and Paro, 2014). Dynamic expression patterns of a number of PRC2-like gene homologs were expressed during cowpea female gametogenesis, including *FIE*, *MSI1*, *EMF1*, *EMF2*, *EMF2*-like, *CLF* and *VRN5* (Fig. 6C). In the mature embryo sac, egg and central cell samples, *FIE*, *MSI1*, *EMF2_2*, *EMF2*-like, and *VRN5_2* were all co-expressed. High levels of *FIE*, *EMF2*-like, and *VRN5_ 2* were evident in the central cell (Fig. 6C). Given the multimeric protein composition of PRC2 complexes, it is not possible to predict the composition of one or more potential complexes that might be functioning during cowpea female gametogenesis from these transcriptional analyses.

### Cell cycle arrest in egg and central cells

Cell cycle genes are relatively well conserved across plant species (Wang et al., 2004, Scofield et al., 2014). Analyses of the expression of cell cycle genes known to block and/or promote passage through various phases of the cell cycle in isolated cell types has been used to predict the stage of cell cycle arrest (Juranic et al., 2018, Sornay et al., 2015, Sukawa and Okamoto, 2018, Velappan et al., 2017, Komaki and Sugimoto, 2012). A core set of cell cycle genes from *Arabidopsis* was used to identify 141 non-redundant cowpea genes encoding proteins with various roles in cell cycle regulation, including 46 cyclins (CYC), 12 cyclin-dependent kinases (CDKs) and 5 CDK inhibitors (interactor/inhibitor of cyclin-dependent kinases/Kip-related proteins; KRP/ICKs) in addition to other key regulators (Supplemental Table 15 and 16). The expression of these in the central cell and egg cell transcriptomes was used to predict the stage of arrest in both cell-types in mature cowpea female gametophytes (Fig. 7).

The transition from G1 to S phase requires activity of CDKA/CYCD complexes to phosphorylate, and inactivate RBR, thereby enabling E2F transcription factors to drive S-phase gene expression (e.g. genes such as *PCNA* and *MCM2-7*; (Dante et al., 2014). Functional activity of the CDKA/CYCD complex at the G1/S transition is also inhibited by KRP/ICKs, which act on additional CDK/CYC complexes to block transition from G2 to M (Dante et al., 2014). The protein kinase, Wee1 can contribute to a block in cell-cycle progression at the G2/M transition (De Schutter et al., 2007). The MYB3R family members, MYB3R1 and MYB3R4 transcriptionally activate many G2/M promoting genes (Haga et al., 2011). MYB3R3 and MYB3R5 negatively regulate transcription of the same genes (Kobayashi et al., 2015). However, MYB3R1 can also negatively regulate transcription, redundantly with MYB3R3 and MYB3R5 at G2/M (Kobayashi et al., 2015).

The egg cell appeared to be arrested in G1/S because expression of RBR and three KRP-like genes *KRP7*/*ICK5_1*, *KRP3*/*ICK6_1* and *KRP3*/*ICK6_2* was moderate to high in the EC. Expression of the *CYCD*s (*CYCD2*, *CYCD3*, *CYCD5* and *CYCD6*), indicative of G1 phase, was low to moderate in the EC while expression of S-Phase genes was moderate to high (Fig. 4, Supplemental Table 16). The high expression of two repressive *MYB3R5* homologs (*MYB3R5_3* and *MYB3R5_4*) and three *Wee1* homologs (*Wee1_1*, *Wee1_2* and *Wee1_3*) in the EC (Fig. 7, Supplemental Table 16) suggests that it is unlikely the EC can progress through the G2/M transition (Kobayashi et al., 2015).

The central cell appeared to have progressed past S phase and appeared to be arrested in G2/M because *RBR1* and *KRP7*/*ICK5_1* expression were strongly reduced compared to that in the EC, as was expression of all S phase-related genes (Fig. 5, Supplemental Table 16). The expression of three *CYCB*-like homologs promoting the G2/M transition, (*CYCB2;3_1*, *CYCB2;3_2* and *CYCB1;2.1_2*) showed high expression in CC at almost three times that observed in the egg cell (Fig. 5, Supplemental Table 16). The CC showed only moderate expression of two G2/M promoting *MYB3R1* homologs, expression of repressive *Wee1_1* and *Wee1_2* was moderate and *MYB3R4* and *MYB3R5* expression was low to undetectable (Fig. 5, Supplemental Table 16). Taken together, these data indicate the cowpea CC is preparing to enter M phase.

### Post-fertilization gene expression in cowpea

The GO terms characterizing the early post-fertilization embryo transcriptome comprised of elongating egg cells, zygotes and early embryos, provided signatures of growth with microtubule movement, cell proliferation, transcription, translation carbohydrate processes, and cell wall biogenesis (Fig. 3A). Meiosis transcript expression was restricted to homologs of *MSH4*, *PRD1*, *OSD1* and *SMG7* (Fig. 3C). A range of auxin biosynthesis-like genes were expressed including *YUCCA*-likes, *2*,*3*,*4*, *6*, *8* and *10* and *TAR10*-like (Fig. 4A). By contrast, a lower repertoire of auxin-responsive type gene expression was observed relative to that seen during male and particularly female gametophyte formation (Fig. 4B). Evidence for extensive microRNA-induced silencing was low given the low levels of *HEN* and *DCL1*. Activity of an *Arabidopsis*-like PTGS pathway was unlikely with many components undetectable (Fig. 5A), and significant RdDM activity at the cell-type stages collected was also doubtful. Expression of other DNA methylation components was evident as was expression of the DNA demethylase *DME* (Fig. 5B). A repertoire of histones genes were expressed and expression of a number of PRC2-like genes suppressed. High levels of *JMJ25*-like gene expression were noticeable suggesting H3K9 demethylation may be supporting early cowpea embryogenesis (Fig. 6D).

## Discussion

### Cowpea reproductive cell-type transcriptomes: a legume resource

The generated cowpea transcriptomes spanning the temporal sequence of events during male and female gametophyte and early seed development enabled detection of enriched, specifically expressed genes including cell-type specific genes expressed at low levels. Verification levels of 71% of cell-type specific expression by qRT-PCR and spatial confirmation *in situ* hybridization confirmed the utility of these transcriptomes for further dissection of regulatory networks in cowpea and syntenic legumes. Comparative analyses of cowpea cell-type specific genes and those specifically expressed in *Arabidopsis*, led to the identification of *Arabidopsis*-derived promoters directing gene expression to meiotic cells, the egg and central cells and early embryos of cowpea. This supports the conservation of some regulatory transcriptional networks during reprodutive cell differentiation in both cowpea and *Arabidopsis*.

The cowpea meiosis genes identified show strong conservation with those found in other angiosperms. Cowpea meiosis genes were, however, found to be expressed at moderate levels in non-meiotic cell-types. *Arabidopsis* tissue RNAseq experiments have also found that meiosis genes are present in many tissues throughout development (Klepikova et al., 2016). This is consistent with their roles in DNA repair, cell-cycle control and recombination, and highlights the likely role of cell-specific protein-protein interactions in determining their molecular function in either meiosis or mitosis (Cromer et al., 2012).

With respect to the involvement of epigenetic pathways during cowpea reproductive development, genes encoding cowpea AGO proteins were identified, and transcriptome analysis suggests an *Arabidopsis*-like RdDM pathway is likely to regulate a number of reproductive stages. *AGO5*, *AGO4* and *AGO6*-like are expressed throughout female gametogenesis. Previous work has suggested that *Arabidopsis AGO5* is expressed in sporophytic tissue and plays an important role in defining the MMC prior to meiosis, but is absent from female gametophyte tissue during subsequent development (Tucker et al., 2012). The roles of RdDM in female gametophyte development may differ between *Arabidopsis* and cowpea, and this can be functionally tested.

By contrast, an *Arabidopsis*-like PTGS pathway does not appear to be functional in the stages analyzed because RDR6 expression, while detectable in leaf, was not expressed in reproductive cell-types. An alternative component may substitute for RDR6 in cowpea reproductive cells. The cowpea cell-type transcriptomes also suggested potential for PRC2-like complex formation and function during male and female gametophyte formation. The data revealed the possibility for a number of novel combinations of proteins in such potential cowpea-PRC2 multimeric complexes. The identified gene candidates could be tested in protein interaction studies, and mutagenic approaches to confirm complex components and their functional relevance. Additional insights into pathways evident during cowpea reproductive development which have been uncovered from *in silico* analyses are discussed further below.

### Plant hormone pathways involved in cowpea reproductive development and patterning

Plant hormones contribute significantly to plant differentiation and growth. In both the megaspore mother cell and late pollen mother cells, GO terms for “abscisic acid (ABA) signalling pathway” and response to abscisic acid were enriched. Underpinning genes were predicted to encode homologs of calcium-dependent protein kinases involved in ABA signal transduction that influence panicle development and seed formation in rice (Ray et al., 2007). Homologues of genes involved in the biosynthesis of brassinosteroids, a HERK receptor-like kinase and a SCARECROW-like transcription factor, involved in perception and positive regulation of brassinosteroid-mediated gene expression, were expressed during female embryo sac mitosis. In addition, homologs of genes functioning in reproductive development in other species, and involved in synthesis and perception of jasmonic acid and cytokinin were upregulated in mature embryo sacs (Li et al., 2004, Bartrina et al., 2011, Kim et al., 2005). Functional roles of these genes in cowpea remain to be determined.

Auxin biosynthesis and auxin-responsive genes were upregulated during early cowpea pollen development, reflecting the functional requirement for auxin in early *Arabidopsis* pollen development (Yao et al., 2018). Auxin biosynthesis is also considered important during *Arabidopsis* female gametophyte development. For example, the auxin biosynthesis genes *YUCCA1* and *YUCCA2* are considered to be required early for cell specification. Later expression of *YUCC8*, *TAA1* and *TAR2* biosynthesis genes are required for nuclear proliferation, vacuole formation and embryo sac growth (Panoli et al., 2015). Studies have suggested an auxin gradient serves as the morphogen driving female gametophyte cell specification in *Arabidopsis* (Pagnussat et al., 2009). However, auxin activity has not been experimentally determined in the developing *Arabidopsis,* maize and *Hieracium pilosella* female gametophytes during mitotic divisions (Lituiev et al., 2013).

While *YUCCA* and *TAR*-like auxin biosynthesis genes were expressed in the cowpea female megaspore mother cell, the expression of auxin biosynthesis genes was barely detectable in subsequent stages of megagametogenesis except for low levels of *TAR2*. Expression of *YUCCA6*, *8* and *10* homologs, and *TAR2* and *TAR4* homologs was detected at low levels in the egg and central cell of the mature embryo sac. The auxin response machinery including multiple *ARF* transcription factors, *AUX-IAA* genes, *SAUR* genes and auxin transporters was detected at moderate to high levels. Highest expression levels were evident in the egg cell and central cell. In the presence of low levels of auxin, AUX-IAA proteins interact with the ARF transcription factors and inhibit transcription of auxin responsive genes. At higher auxin levels, AUX-IAA proteins are degraded by the ubiquitination pathway, freeing the ARF transcription factors to promote expression of their target genes (Wang and Estelle, 2014).

Given that expression of auxin biosynthesis genes was very low in the developing cowpea female gametophytes, this may indicate that some auxin-related processes are suppressed. The contrasting pattern of very low levels of auxin biosynthesis gene expression and higher relative abundance of auxin responsive genes and transporters suggests that if auxin is involved in cowpea female gametophyte patterning it may be transported in from the sporophytic tissues post-meiosis. Experiments using auxin sensors in transgenic cowpea may resolve this possibility. Alternatively, auxin may have effects in cowpea sporophytic tissue that indirectly affect cell fate decisions in the female gametophyte as has been suggested for *Arabidopsis* and maize (Lituiev et al., 2013).

### Cell cycle arrest in male and female cowpea gametophytes: involvement of epigenetic regulation

Unlike most other eukaryotes, where the gametes remain in the G1 cell cycle phase through nuclear fusion at fertilization (karyogamy), higher plants form gametes that undergo karyogamy in G1, S or G2 phases of the cell cycle (Friedman, 1999). We observed that cowpea pollen is predominantly bi-cellular, containing a vegetative and a generative cell at maturity, as approximately 10% of pollen grains contained a vegetative cell and two sperm cells. Therefore, the events of pollen mitosis II in cowpea continue after pollen-tube germination. *Arabidopsis* pollen grains are tri-cellular and sperm cells progress through S-phase after pollen germination to arrive at G2 immediately prior to fertilization (Friedman, 1999).

Although the cell cycle stage of cowpea sperm cells at fertilization remain to be determined, similarities in histone transcription were observed between *Arabidopsis* and cowpea during pollen maturation. Histone *H3.3* and *H3.1* transcripts were detected during cowpea male gametophyte formation, but histone *H3.1* transcripts were not detected in the mature cowpea pollen-enriched sperm cell sample, as is also observed in *Arabidopsis*. H3.3 variants correlate with gene expression in *Arabidopsis*, while H3.1 variants are incorporated during the S-phase of the cell cycle (Borg and Berger, 2015), and references therein). In rice, EMF2 function is essential for pollen development, and it is thought to interact with a rice MSI1 homolog, PTC1 (Deng et al., 2017). Genes encoding homologs of a number of these PRC2 components were also expressed in cowpea pollen development and may have comparable roles.

The central cell and egg cell are arrested at different stages of the cell-cycle prior to fertilization. Studies in maize, rice, barley and *Arabidopsis* suggest that the egg is arrested at G1 (Chen et al., 2017, Mogensen and Holm, 1995, Sukawa and Okamoto, 2018), while, in *Arabidopsis,* it has been proposed that the central cell is most likely to be at G2 (Berger et al., 2008). Analysis of cell cycle gene expression in cowpea egg and central cells indicated their likely arrest in G1/S and G2/M cell cycle phases, respectively, consistent with that for *Arabidopsis*.

Epigenetic pathways involving RdDM and DNA methylation repress transposon activity and regulate female gametophyte arrest. DNA methylation is very low in the central cell and reduced in the egg cell (Kawashima and Berger, 2014). RdDM and DNA methylation are decreased in the central cell, and this allows expression of transposons, which are normally repressed. Subsequent production of siRNAs that then direct methylation within the egg cell, prevent transposon activation (Ibarra et al., 2012). In cowpea, expression of *AGO4*, *AGO5* and *MET1* homologs was several times lower in the central cell than in the EC, suggesting that *de novo* RdDM may also be less active in the central cell. Furthermore, the miRNA pathway is likely more active in the cowpea egg cell than in the central cell as the key components, *AGO1* and *DCL1* are expressed around five times higher in the egg cell. This corresponds with observations of Wuest et al. (2010), who reported high egg cell expression levels of *DCL1* and *AGO1* in *Arabidopsis*.

The transcriptome analyses also suggested that one or more novel PRC2-like complexes may function to repress female gametophyte initiation in the absence of fertilization via histone methylation. The conservation of a post-fertilization role of auxin in facilitating embryo patterning in cowpea is supported by the observation of increased transcription of auxin biosynthesis genes in the embryo as found in other species (Hands et al., 2016).

## Conclusions

In this study, we have isolated and analysed cell-specific transcriptomes spanning the events of male and female gametogenesis and early seed formation in cowpea. Cell-specific genes were identified and validated using a combination of techniques including qRT-PCR and *in situ* hybridization. The transcriptomes have identified gene expression pathways in common with reproduction in other angiosperms, in addition to novel differences that can be functionally examined. These cowpea reproductive cell-type transcriptomes provide a useful resource for the further isolation of cell-type specific promoters, and uncovering informative pathways which may aid in improving reproductive resilience and yield in cowpea and related legumes.

## Materials and Methods

### Plant material, tissue collection and generation of RNA sequences

Cowpea plants (*Vigna unguiculata* IT-86D-1010) were grown in the glasshouse under previously described conditions (Salinas-Gamboa et al., 2016). To collect RNA samples, tissues at appropriate developmental stages were fixed and processed for LCM according to previously published protocols (Okada et al., 2013). To collect sperm cells, anthers and stigmas were collected from flowers at anthesis and subjected to osmotic shock in Brewbaker and Kwack medium (Brewbaker and Kwack, 1963), pH 6.5, supplemented with 12.5% (w/v) sucrose in 15 ml centrifuge tubes. The homogenate was centrifuged at 130 rpm, for 30 minutes at room temperature, then filtered through 150 µm and then 30 µm CellTrics nylon sieves (Partec GmbH) and collected in 2 ml microfuge tubes. The homogenate was centrifuged at maximum speed for 2 minutes to pellet cells, and 50 µl was layered on 0.5ml of 10% Percoll in 2ml microfuge tubes that were centrifuged at 900xg for 2 minutes. 1.5 ml of a solution of 0.52 M (10%) mannitol, 10 mM MOPS buffer (pH 7.5) was added, and the tubes were centrifuged for 2 minutes at maximum speed. 20µl of the resulting pellet was added to 30ul of ARCTURUS PicoPure RNA isolation buffer and snap frozen in liquid nitrogen before storing at −80C. Total RNA was extracted from the LCM samples and sperm cell enriched samples using the ARCTURUS® PicoPure® RNA Isolation Kit (Applied Biosystems®) according to the manufacturer’s protocol. RNA was subjected to two rounds of amplification (three rounds for fTET, ES2n and ES4n) using the MessageAmp II RNA amplification kit (Ambion). Total RNA was extracted from leaf as described (Spriggs et al., 2018).

### Sequencing and read alignment

Illumina sequencing libraries were prepared from the RNA samples by the Australian Genome Research Facility (AGRF) and run on the Illumina HiSeq2500 sequencing platform multiplexed in 3 lanes. At least 36 million reads were produced per sample (approximately 1.6 billion reads, total) of which ∼86% could be uniquely aligned to the IT86D-1010 genome (Spriggs et al., 2018) using the BioKanga software package (Supplemental Table 1, Supplemental Data 8). A total of 74,839 genes were predicted in the cowpea IT86D-1010 genome (Spriggs et al., 2018) using Augustus v 3.1.0 (Stanke and Waack, 2003) and *Arabidopsis thaliana* (TAIR 10) as the gene training set. As no correlation between gene length and number of reads aligned was observed (Pearsońs correlation coefficient = - 0.010), the Reads-Per-Million (RPM) normalization factor was used in all cases reporting normalized read counts. The RPM normalized counts between replicates of a cell-type sample were strongly positively correlated, with a Pearson’s correlation coefficient (PCC) over 0.8 for most samples. Exceptions were replicates in the megaspore mother cell (MMC; PCC = 0.72), two-nucleate female gametophyte (ES2n; PCC= 0.766) and the pollen microspore (MIC; PCC = 0.62; Supplemental Fig. 1).

### R versions and rsgcc analyses

Analyses made using R software packages used R Studio version 1.0.143, with R version 3.4.3 running under Windows 10. The rsgcc software of Ma and Wang (2012) was used as described by Zhan et al. (2015). The mean RPM normalized counts from each cell-type was used together with the getsgene() function of the rsgcc R package (rsgcc version 1.0.6) with the following parameters “getsgene(data, Log = FALSE, tsThreshold = 0.50, MeanOrMax = “Max”)”.

### Gene Ontology annotations

Annotations were made by aligning the predicted IT86D-1010 coding sequences (Supplemental Data 9) against the NCBI nr protein database (version December 23^rd^ 2017) using the DIAMOND aligner (Buchfink et al., 2015), version 0.9.13 returning at most ten alignments with a maximum e-value of 1e-5. The generated XML file was supplied to Blast2GO (version 5.0.13 running on windows 10 with Java version 1.0.8 152), which was run with default parameters, to assign Gene Ontology (GO) annotations (Ashburner et al., 2000, The Gene Ontology, 2017) to each predicted gene. This approach assigned putative functional annotations to 54,223 predicted genes, with 39,653 predicted genes assigned GO terms (Supplemental Data 1).

### Filtering to identify cell-specific genes in the egg cell, central cell and sperm cell transcriptomes

Genes specifically expressed in central cell, egg cell and sperm were selected using the filter function in Microsoft Excel to choose genes with a mean RPM at least five times higher in the cell-type of interest than any other cell-type (see full criteria in Supplemental Table 2). No filter was applied to other cell-types in the female gametophyte when selecting egg cell and central cell genes where the contrast between those two cells was the key criteria.

### Differential expression analysis

Differential expression analysis was carried out using the DESeq2 (version 1.18.1) software package in R (Love et al., 2014). Analyses were carried out as described except that differentially expressed genes in each cell-type were identified using the contrast argument of the results function with parameters set to contrast the expression of each gene in one cell-type with the average expression of all other cell-types “results(dds, contrast = list(c(cell_X), c(all_other_cells)), listValues = c(1, −1/13), alpha = .05)”. Genes with adjusted p value < 0.05 and log2 fold change > 2 were selected as significant (Supplemental Data 5).

### Principal Component Analysis

Principal component analysis was carried out on each replicate of our developed transcriptomes using the base R prcomp() function on read counts transformed by the variance stabilising transformation, vsd(), function of DESeq2. Principal components were plotted in 3D using the plot3d() function of the rgl package. Colours suitable for all vision types were chosen from the colour alphabet previously described (Green-Armytage, 2010). Plots were exported as .svg files and figures were assembled in Adobe Illustrator.

### Comparisons between cowpea and Arabidopsis female gametophyte LCM data sets

The best *Arabidopsis* homolog for each predicted cowpea gene was identified by using the DIAMOND aligner (Buchfink et al., 2015) (version 0.9.13) to align cowpea genes against the TAIR 10 gene database, and selecting the best match for each cowpea gene (Supplemental Data 7). *Arabidopsis* gene sets from the MMC (Schmidt et al., 2011), ovule nucellus, early mitotic female gametophytes (Tucker et al., 2012) and the egg and central cell (Wuest et al., 2010, Steffen et al., 2007) were compared with the *Arabidopsis* annotations of each cowpea gene in the differentially expressed genes (log2 fold-change >2) in corresponding cell-types from the cowpea dataset.

### Identification of core meiosis, auxin biosynthesis, auxin-responsive genes and cowpea cell-cycle genes

A set of 18 core cowpea meiosis genes were identified by BlastP search of key genes in the literature (Supplemental Table 7 and references therein) against the amino acid sequence of the cowpea IT86D-1010 predicted genes. In each case the best match based on bit-score was chosen for further analysis. To identify cell-cycle related cowpea genes, the amino acid sequences of a set of cell-cycle genes from *Arabidopsis* (Juranic et al., 2018) were used in BlastP searches against the predicted cowpea protein sequences. BlastP results with an e value < 1e-20 were chosen, and the annotation of cowpea sequences matching multiple *Arabidopsis* genes was chosen based on the highest bit score. The *KRP* gene family and *cdc*25 homologs were selected with an e value < 1e-10. Where multiple cowpea genes matched a single *Arabidopsis* gene, these were notated with an underscore and number after the gene name starting with 1 for the best match and increasing (e.g. VuGene_1). Auxin-related genes were identified in the laser captured cell-type transcriptomes, based on their GO annotations (Supplemental Table 9).

### Identification of cowpea RNA silencing, RdDM and histone genes

A total of 37 key genes in RNA silencing and RdDM pathways were identified by searching the annotations produced by the Blast2GO software for genes described in key literature (Supplemental Table 10). Fragments not encoding full length protein homologs were not included in further analysis.

A total of 49 putative histone encoding genes and 20 JMJ homologs were identified in annotations produced by Blast2GO as described above. The predicted histone variant encoded by each gene was assigned based on annotations, and on homology of the predicted protein to *Arabidopsis* histone amino acid sequence (Supplemental Fig. 7).

### Phylogenetic tree construction

Phylogenetic trees were made using Geneious version 10.2 (Biomatters, https://www.geneious.com). Trees were built using the following parameters: ClustalW multiple-sequence alignment with BLOSUM cost matrix, Gap open cost 10, Gap extend cost 0.1, Jukes-Cantor Neighbour-Joining consensus tree, no outgroup, Bootstrap resampling with 1000 replicates, 50% support threshold.

### Validation by qPCR

Whole tissues were collected from the following tissues, emerging leaves, developing stem, petiole, shoot apical meristems and roots (Supplemental Table 3), frozen in liquid nitrogen and RNA was extracted with a QIAGEN RNeasy Plant mini kit according to the manufacturer’s protocol, including on-column DNAse treatment. cDNA was synthesised using 1ug of RNA with the SuperScript™ III First-Strand Synthesis System (Thermo Fisher Scientific). Central cell and sperm genes were selected for testing by qPCR and *in situ* hybridisation based on specificity in expression and putative function of the predicted genes. Egg cell genes were chosen for testing based on the specificity of their expression and homology to *Arabidopsis* genes known to be specifically expressed in egg cell. Genes in other cell-types tested were selected based on their specificity score in the rsgcc analysis and on our ability to design primer pairs (Supplemental Table 17) that specifically amplified the predicted region, including intron structures, from genomic and cDNA. qRT-PCR was carried out on a Roche Light Cycler ® 480 thermal cycler using SYBR Green I master mix according to the manufacturer’s protocols. Quantification was done using LinRegPCR software as described (Ruijter et al., 2009, Ramakers et al., 2003).

### In situ hybridization

ISH was performed as previously described (Jackson, 1992) with some modifications for improved resolution in reproductive organs of cowpea. For ISH performed on sectioned specimens, mature unpollinated ovules were fixed in 4% paraformaldehyde and embedded in Paraplast. Sections at 12 μg thickness were attached to ProbeOnPlus slides (Fisher Biotech) and processed as previously described (Jackson, 1992). For whole-mount ISH, developing ovule primordia, unpollinated mature ovules, and mature anthers were fixed in paraformaldehyde (4% paraformaldehyde, 2% Triton, and 1× PBS in diethylpyrocarbonate DEPC-treated water) for 2 h at room temperature with gentle agitation, washed three times in 1× PBS-DEPC water, and embedded in 15% acrylamide:bisacrylamide (29:1) using pre-charged slides (Fisher Probe-On) treated with poly-l-Lys as described (Bass et al., 1997). Fixed pollen grains were stripped from the anthers before embedding, and incubated with an enzymatic solution containing 1% driselase, 0.5% cellulose, and 1% pectolyase, for 30 minutes (min) in a humid chamber at 37°C. All samples were subsequently incubated with 0.2M HCl solution for 30 min at room temperature (RT) and in 1ug/ul proteinase K for 30 min at 37°C, before being processed as previously described (García-Aguilar et al., 2005). Slides containing paraffin sections were mounted on Cytoseal, whereas slides containing whole specimens were mounted in either 50% (ovules) or 20% (pollen grains) glycerol, and analyzed with a Leica DMRB microscope under Nomarski illumination.

### Plasmid construction and cowpea transformation

Gateway entry vectors containing cell-type specific promoter elements together with genes encoding fluorescent proteins were gifts from Shai Lawit and Marc Albertsen (Lawit et al., 2013) as follows: *AtDD1_pro_:ZsYellow1*, *AtDD9_pro_:DsRed-Express*, *AtDD25_pro_:DsRed-Express*, *AtDD31_pro_:AcGFP1*, *AtDD45_pro_:DsRed-Express*, *AtDD65_pro_:AmCyan1*, *AtLat52_pro_:AcGFP1*, *AtRKD2_pro_:DsRed-Express*. The promoter element of *AtEC1.1* was amplified from *Arabidopsis* Col-0 DNA by PCR with primers including adapter elements, and cloned into gateway entry vectors containing AmCyan1. The *AtDUO1* promoter was synthesized and cloned into pMK-RQ by GeneArt (Thermo Fisher Scientific) with SfiI and BglII sites introduced for subsequent cloning into gateway entry vector containing AcGFP1.

A gateway-compatible binary vector pOREOSAr4r3 was created by insertion of a Gateway recombinational cassette amplified from pDESTr4r3 (Invitrogen) into ClaI and KpnI sites of pOREOSA vector backbone, which was a gift from Thomas J. Higgins, CSIRO. Reporter entry clones were introduced into the pOREOSAgw expression vector via multisite Gateway recombination. Final constructs contained either triple or quadruple fluorescent labels (Supplemental Table 20). AtKNU:YFP:3’KNU cassette was previously described (Tucker et al., 2012) and was inserted into the pOREOSA via HindIII digestion. Gateway cloning reactions were performed with LR Clonase II Enzyme mix (ThermoFisher Scientific). All plasmid vectors were verified by sequencing and final constructs electroporated into *Agrobacterium tumefaciens* strain AGL1 for use in cowpea stable transformation (see below). Details of the primer sequences and constructs transformed into cowpea are described in Supplemental Tables 19 and 20. Cowpea transformation was carried out using *Agrobacterium*-mediated transformation of cotyledonary nodes (Popelka et al., 2006). Transformed explants were first selected on a medium containing 100 mg/L kanamycin and then on 20 mg/L geneticin (G-418) for up to 3-6 months. Shoots developing healthy roots were transferred into 90mm small pots containing sterilized soil mixture (Van Schaik’s Bio-Gro Pty Ltd, Australia), acclimatized in the growth room at 22°C with 16h photoperiod for up to 4 weeks, and then transferred to the glasshouse in larger 4.5L pots. PCR was performed to confirm the presence of the *nptII* and reporter genes with the primers listed in Supplemental Table 19. The number of independent cowpea transgenic lines per construct varied from three to 15.

### Fluorescent microscopy and clearing procedures

Samples were examined using a Zeiss fluorescence microscope (Axio Imager M2; Carl Zeiss) with an HXP 120 light source. Digital images were captured using an AxioCam HRc camera and ZEN 2.6 software (Carl Zeiss). Signals from dsRED-Express were observed with the Zeiss filter set 43, AcGFP1 signals were observed with the Zeiss filter set 13, AmCyan1 signals were observed with Zeiss filter set 47, while YFP and ZsYellow1 signals were observed with Zeiss filter set 46. The exposure time was adjusted to as appropriate for each sample. Images were processed in ZEN2.6 software and assembled in Adobe Photoshop. The method previously described in (Salinas-Gamboa et al., 2016) was used when tissue clearing was required. For light microscopy, the Zeiss Axioskop 2 microscope equipped with Nomarski optics, Spot Flex colour camera and Spot 5.1 software was used to capture images (Diagnostic Instruments, Inc).

### Generation of transgenic *Arabidopsis* with cowpea promoter constructs

The regulatory region of eleven cowpea genes was transcriptionally fused to the *uidA* (*GUS*) reporter gene by amplifying a fragment corresponding to either the complete intergenic region (*VuNB-RCpro; VuSDR1pro*) or an arbitrary fragment (*VuBAS1pro; VuGULL06pro; VuRKD1pro; VuEC1.1pro; VuGIM2pro; VuXTH6pro; VuXTH32pro; VuAt3g09950pro; VuKNU-L1pro*) located upstream of its coding sequence but excluding the 5 untranslated region (see Suppl Table 21 for primers). Amplicons were cloned into pCR8 TOPO TA (Invitrogen) and used as donors in LR recombination (LR Clonase II; Invitrogen) with pMDC162 (Curtis and Grossniklaus, 2003), producing a binary vector that contains the *uidA* reporter gene. Transgenic Col-0 plants were obtained by floral dipping as previously described (Clough and Bent, 1998). At least 15 T1 individuals were obtained and analyzed to quantify the frequency of GUS expression at different stages of male and female reproductive development. GUS staining assays for stages before fertilization were conducted as described (Vielle-Calzada et al., 2000).

## Accession numbers

**Large datasets**

.bam files of sequences from TO BE CONFIRMED

Fasta.gz archives of raw reads for MMC, EM and Sp

## Supplemental Material

**Supplemental Tables Supplemental Table 1. Align stats**

Number and proportion of Illumina reads aligned against the IT86D genome for each sample

**Supplemental Table 2. Filtering**

Criteria and results of selecting genes based on filtering in Excel

**Supplemental Table 3. Arabidopsis Reproductive genes and promoters**

Summary of *Arabidopsis* reproductive cell-specific gene expression of their Cowpea homologs in cowpea and activity of their promoters driving transgenes in *Arabidopsis*

**Supplemental Table 4. qPCR**

Table of tissues and results of qRT-PCR validation of cell-specific gene expression

**Supplemental Table 5. CS_DE_gene_overlap**

Table summarising the number of common genes between cell-specific and differentially expressed datasets.

**Supplemental Table 6. Compare datasets**

Comparison of published cell-specific datasets with our transcriptomes

**Supplemental Table 7. Meiosis genes**

Names and references for the cowpea homologs of genes involved in meiosis

**Supplemental Table 8. Meiosis RPM**

Expression of meiosis genes as RPM counts

**Supplemental Table 9. Auxin genes**

Expression of annotated auxin related genes as RPM counts

**Supplemental Table 10. RNAi, RdDM refs**

Names, functions and references for the cowpea homologs of genes involved in RNA silencing and RdDM

**Supplemental Table 11. RNAi, RdDM RPM counts**

Expression of RNAi and RdDM genes as RPM counts

**Supplemental Table 12. Histone RPM counts**

Expression of histone genes as RPM counts

**Supplemental Table 13. PRC RPM counts**

Expression of PRC genes as RPM counts

**Supplemental Table 14. JMJ RPM counts**

Expression of JMJ genes as RPM counts

**Supplemental Table 15. Cell-cycle genes**

Names and references for the cowpea homologs of genes involved in cell-cycle progression

**Supplemental Table 16. Cell-cycle RPM**

Expression of cell-cycle genes as RPM counts

**Supplemental Table 17. Oligonucleotide sequences for qRT-PCR**

Sequences of oligonucleotide primer sequences used for qRT-PCR in this study

**Supplemental Table 18. Probe sequences**

Sequences used in RNA *in situ* probes

**Supplemental Table 19. Cloning Primers**

Sequences of primers used in creating plasmids used in this study

**Supplementary Table 20. Plasmid constructs**

List of plasmid constructs used in this study

**Supplemental Table 21. Activity of Cowpea promoters**

Activity of cowpea promoters in *Arabidopsis thaliana*

## Supplemental Data

**Supplemental Data 1: IT86D-1010 genes and GO annotation**

Large table with GO annotations and descriptions for all cowpea predicted genes

**Supplemental Data 2: Mean RPM counts**

Large table with the average RPM counts for all genes in all cell-types

**Supplemental Data 3: Cell-specificity scores and RPM counts**

Large set of tables with all genes in each of the cell-specific gene sets

**Supplemental Data 4: Cell-specific gene annotations**

Large set of tables with all cell-specific genes for each cell-type and their GO annotations

**Supplemental Data 5: log2 FC2 DE genes**

Large set of tables with all genes DE at log2 fold-change 2 in each of the cell-types

**Supplemental Data 6: DE enriched GO terms**

Large set of tables with the enriched biological process GO terms from each of DE gene sets

**Supplemental Data 7: BlastP comparing Vu and At**

Large table containing the best *Arabidopsis* protein match by homology to the translated coding sequence of each predicted cowpea gene

**Supplemental Data 8: Raw aligned read counts**

A csv file with the raw counts of aligned to each gene in each replicate.

**Supplemental Data 9: Predicted gene coding sequences**

A fasta file with the coding sequences of all IT86D-1010 predicted genes.

## Acknowledgments

We are grateful to Tracy How and Dilrukshi Nagahatenna for their assistance with cowpea transformation and growth, to Natalia Bazanova for assistance with cloning, and to Steven Henderson for suggestions concerning the *in silico* identification of cowpea cell-specific genes.

---

**Supplemental Figure. 1.**
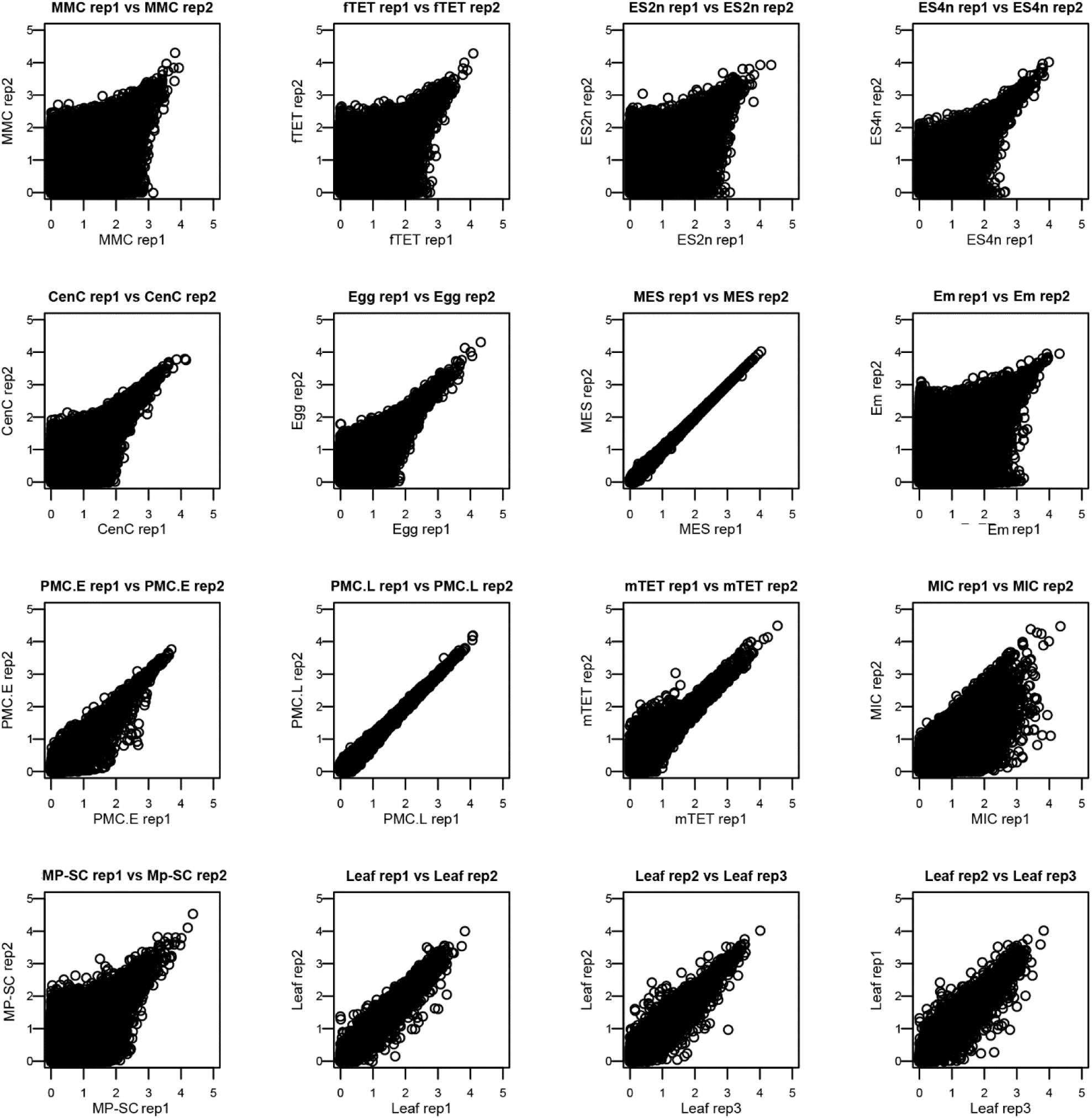
Correlation between RNAseq replicates. Scatter plot of log-transformed mean RPM counts of replicates for each tissue. Replicate 1 and replicate 2 for each cell-type are shown on the Y and X axes, respectively.

**Supplemental Figure. 2.**
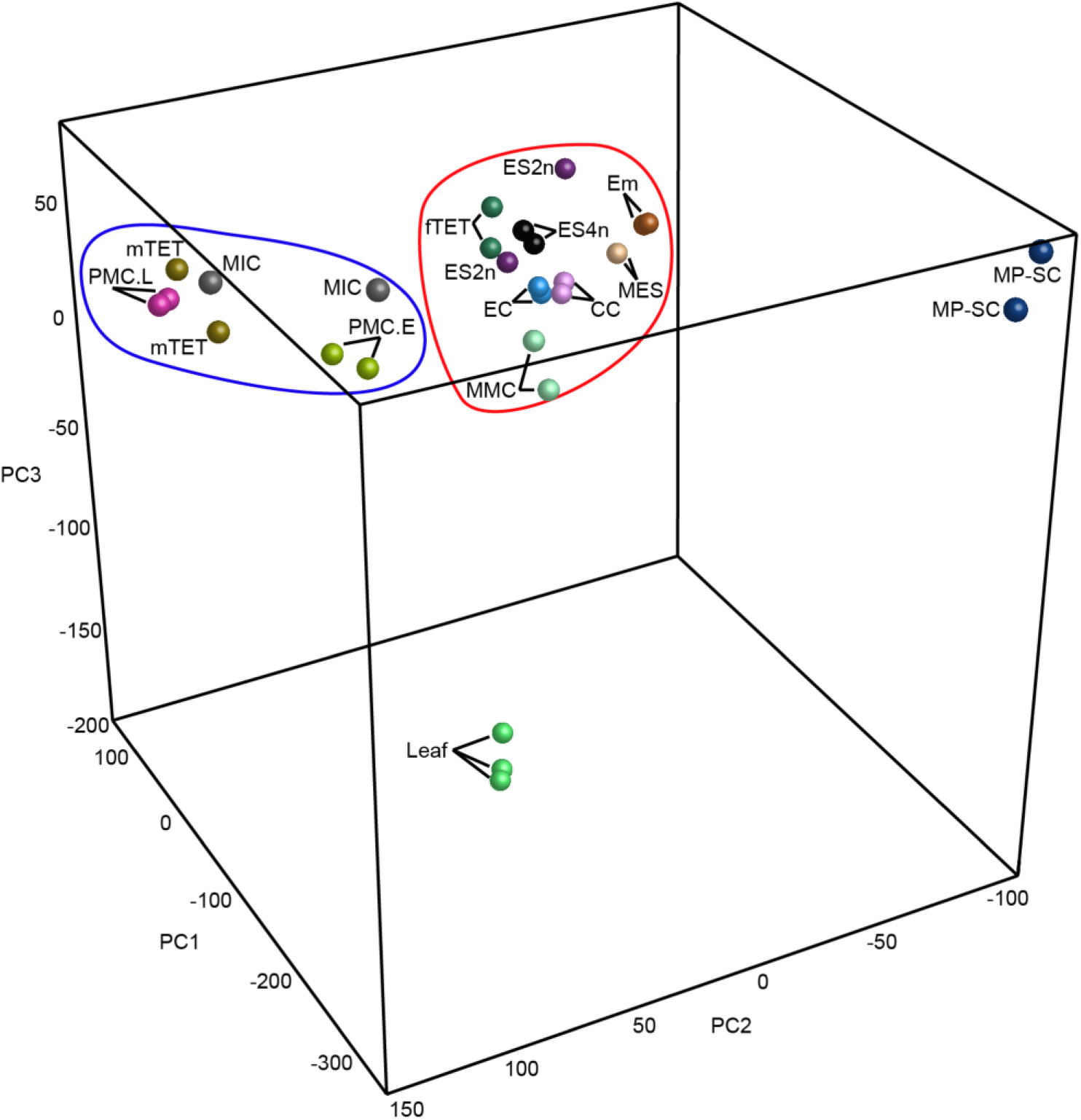
Principal component analysis (PCA). PCA plot showing relationships between all transcriptomes. Abbreviations: MMC, megaspore mother cell; fTET, female tetrads; ES2n, mitotic embryo sac with 2 nuclei; ES4n, embryo sac with 4 nuclei; MES, mature embryo sac at anthesis, containing the egg cell (EC) and central cell (CC); Em, embryo; PMC.E, early pollen mother cell; PMC.L, late pollen mother cell; mTET, male tetrads; MIC, uninucleate microspore; MP-SC, mature pollen-sperm cell.

**Supplemental Figure. 3.**
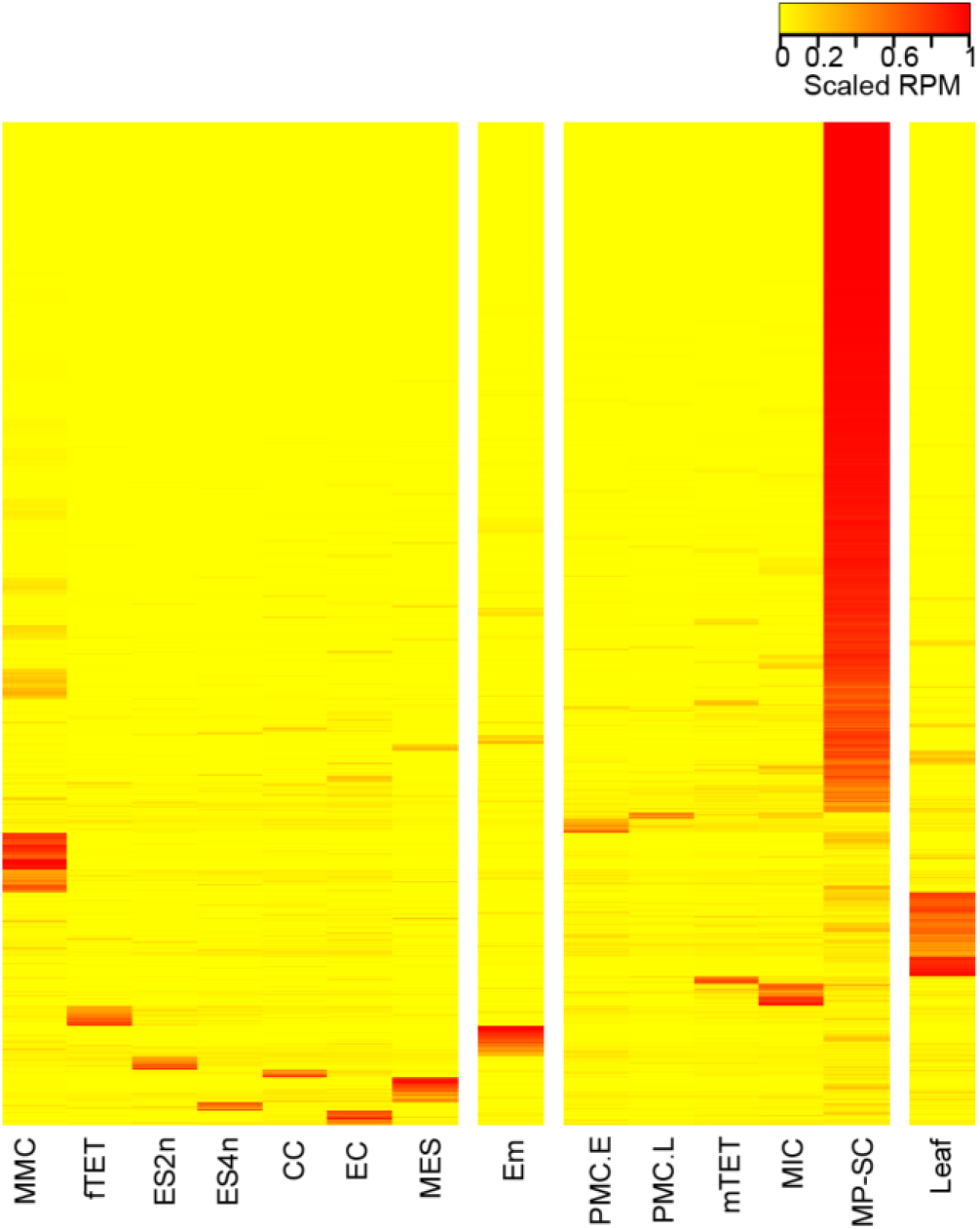
Cell-specific genes expression Heat-map of scaled RPM expression values of the cell-specific genes identified using the rsgcc software package

**Supplemental Figure. 4.**
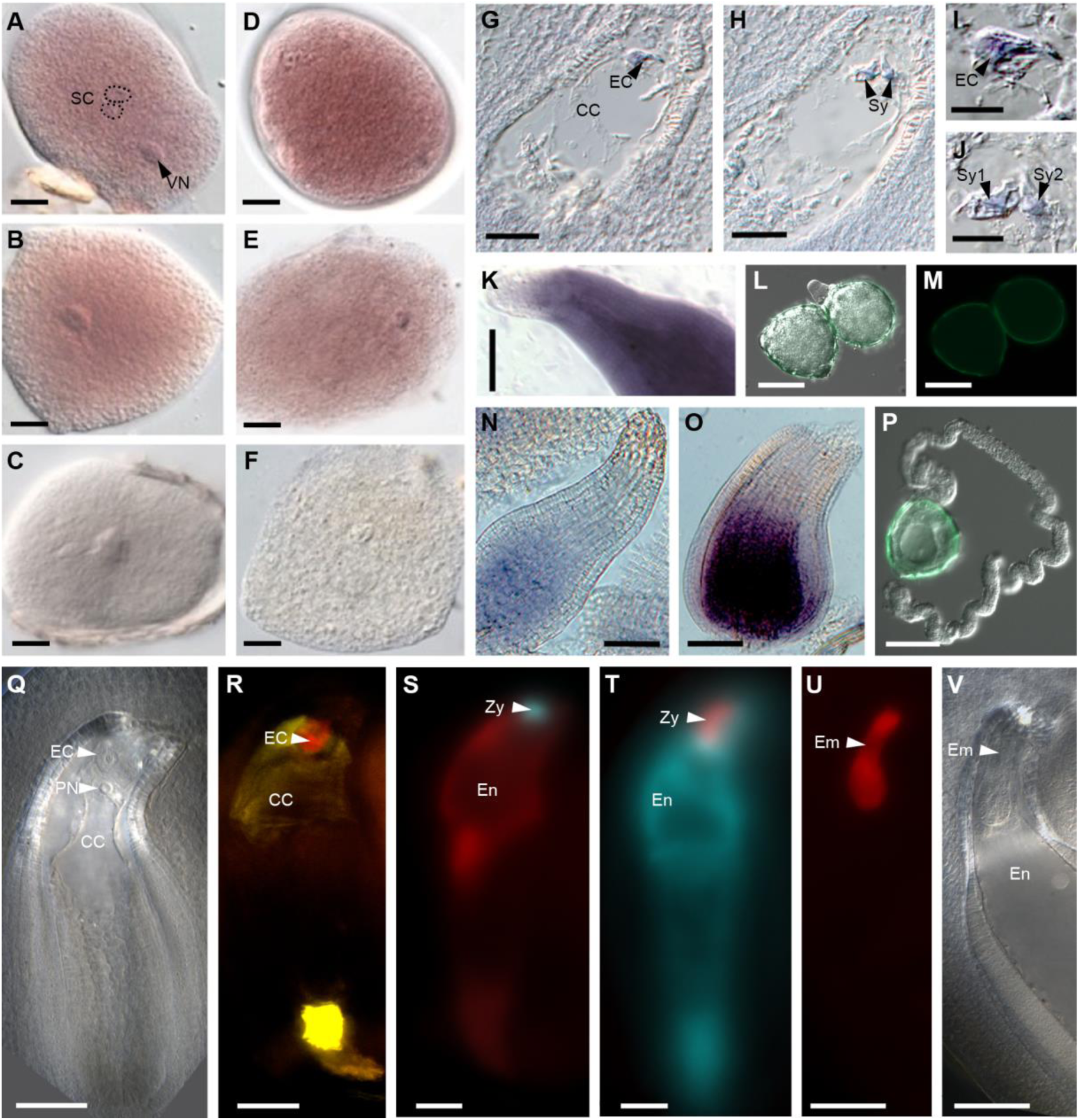
*In situ* hybridization of cell-specific cowpea genes in gametophytic tissue and fluorescent reporters expressed from *Arabidopsis* cell-specific promoters in cowpea. A-B, *VuXTH32* mRNA localized in the sperm cells and vegetative cell cytoplasm. C, *VuXTH32* sense control. D-E, *VuAt3g09950* mRNA abundantly localized in the sperm and vegetative cells. F, sense *VuAt3g09950* control. G, Paraffin section through a fully differentiated female gametophyte showing *VuEC1.1* mRNA localization in the egg cell. H, Consecutive section to (G) showing *VuEC1.1* mRNA localization in the synergids. I, Detail of G showing mRNA localization in the egg cell. J, Detail of H showing mRNA localization in the synergids. K, Whole-mounted ovule showing *VuMEE23* mRNA localization in the central cell and adjacent nucellar cells. L-M, *AtDUO1_pro_:AcGFP1* no expression. N, Whole-mounted ovule showing *VuSDR1* mRNA localization in the central cell. O, Whole-mounted ovule showing *VuGULL06* mRNA localization confined to the central cell. P, No expression of *AtLAT52_pro_:AcGFP1* in germinating pollen. Q, Mature female gametophyte with egg cell, polar nuclei and central cell in cleared ovule. R, *AtDD45_pro_*:*dsRED-Express* (red) in EC and general pattern of the *AtDD1_pro_*:*ZsYellow1* (yellow) in cells external to the embryo sac and in CC. S, *AtEC1.1_pro_:AmCyan1* (cyan) in zygote and *AtDD25_pro_:dsRED-Express* (red) in early endosperm. T, *AtRKD2_pro_*:*dsRED-Express* (red) in zygote and *AtDD9_pro_:AmCyan1* (cyan) in early endosperm. U, *AtRKD2_pro_*:*dsRED-Express* in early embryo. *AtDD9* is not expressed at this stage. V, Cleared ovule showing early embryo corresponding to the stage in U. Scale bars: A-F, I-J = 10 µm; G-H = 18 µm; K, N-O = 35 µm; L-M, P-T = 50 µm; U-V = 100 µm Abbreviations: EC, egg cell; CC, central cell; Sy, synergids; SC, sperm cells; VN, vegetative nucleus; PN, polar nuclei; Em, embryo; En, endosperm; Zy, zygote.

**Supplemental Figure. 5.**
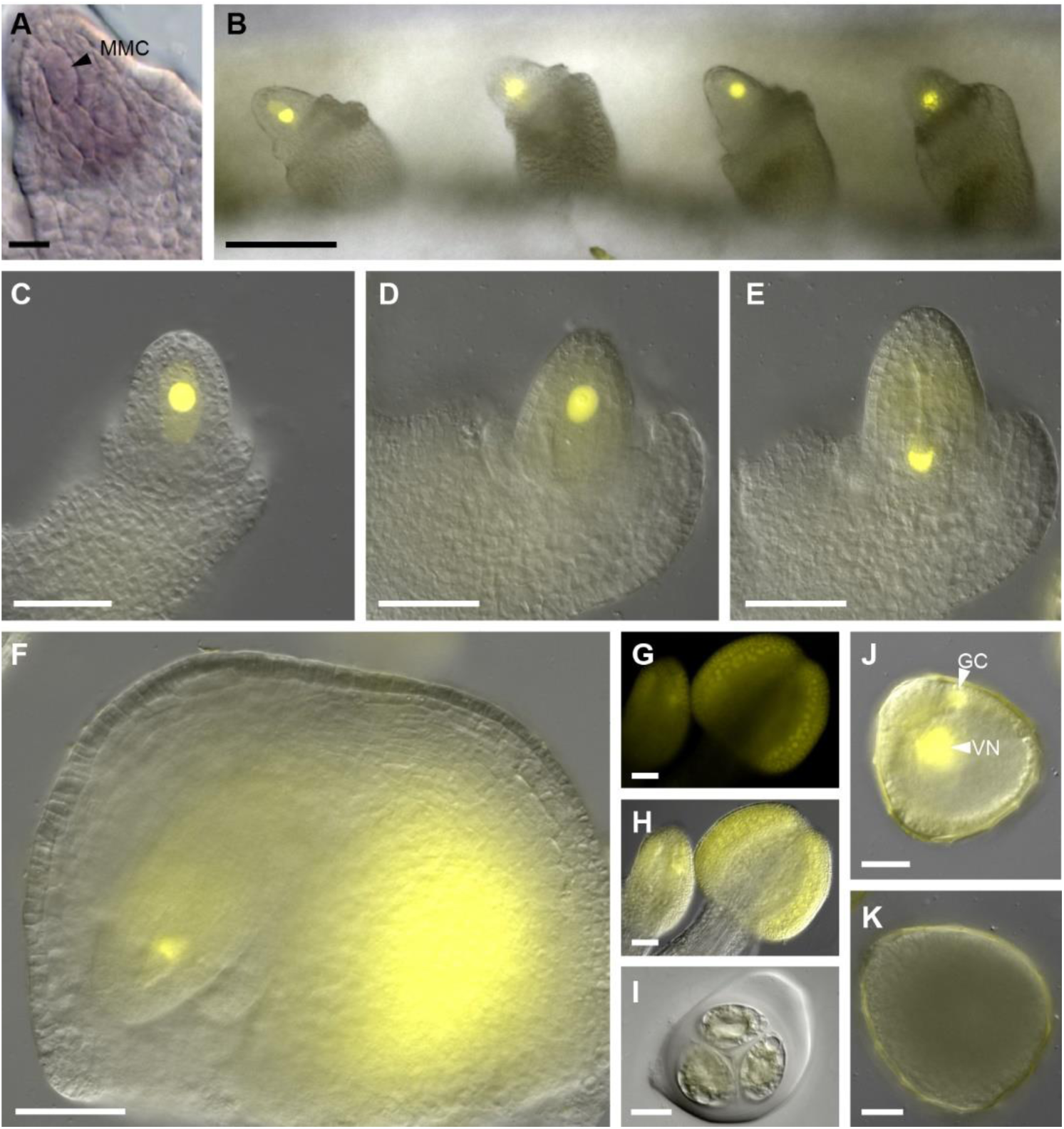
Expression of the KNU genes in cowpea. A, Whole-mount pre-meiotic ovule showing *VuKNU1* mRNA localized in the MMC and the developing nucellus at the onset of integument initiation. B-D, Micrographs of cowpea ovules expressing a *AtKNU_pro_:nlsYFP* reporter construct marking MMC fate. E-F, At the end of meiosis the KNU promoter turns off in female functional megaspore, but the expression persists in somatic ovule cells at the chalazal end. G-H, Expression of *AtKNU_pro_:nlsYFP* reporter construct during male meiosis. I, Male tetrad showing no expression of reporter construct. J, After the first pollen mitosis, the expression of reporter construct is evident in both vegetative and generative nucleus. K, Mature pollen grain does not show any expression. Abbreviations: MMC, megaspore mother cell; GC, generative cell; VN, vegetative nucleus. Scale bars: A = 10 µm; B = 100 µm; C-H = 50 µm; I-K = 20 µm.

**Supplemental Figure. 6.**
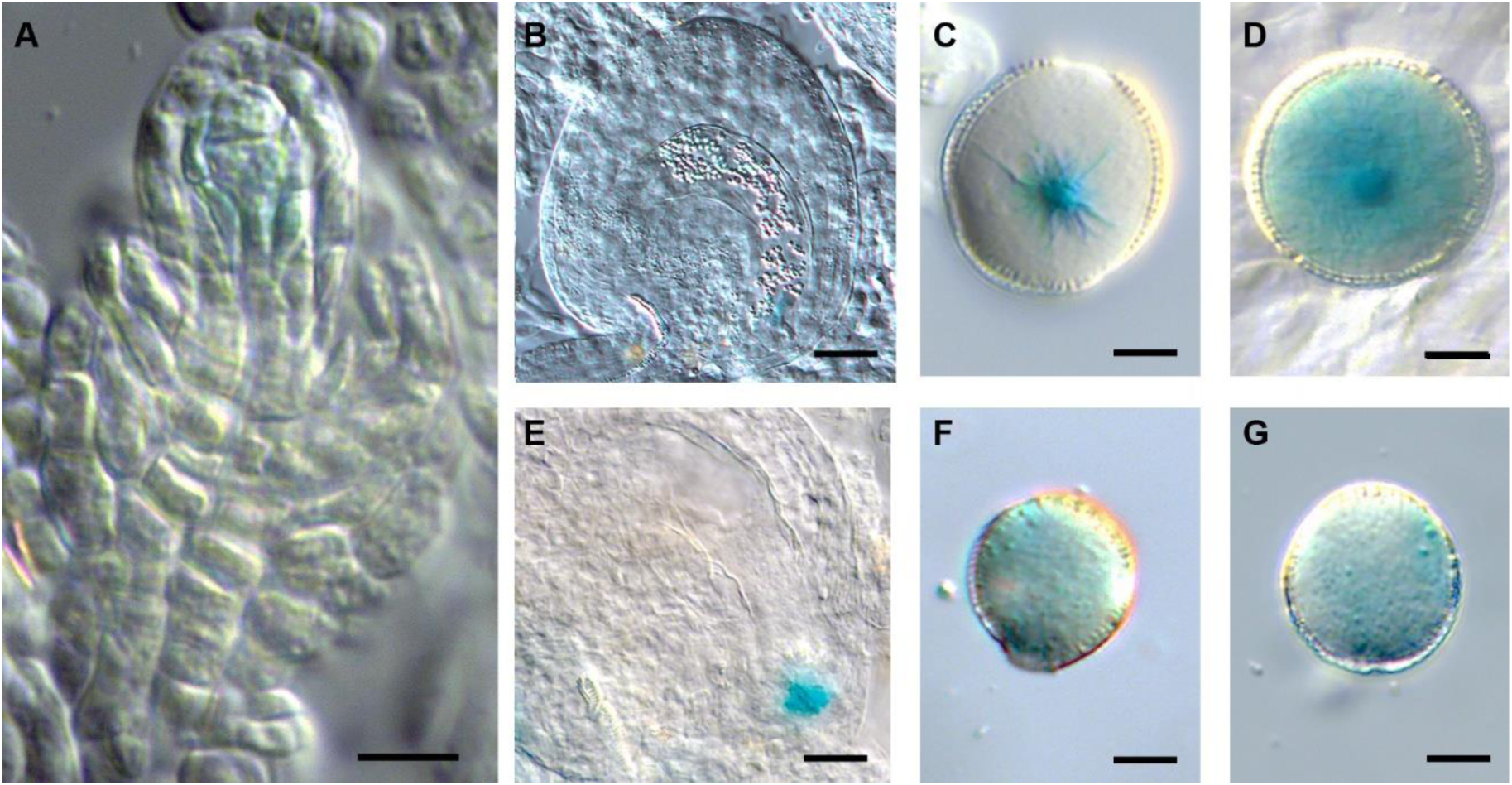
Specific activity of cowpea promoters in reproductive cells of *Arabidopsis thaliana*. A, *VuKNU-L1_pro_:GUS* expression in the developing ovule of *Arabidopsis*. B, *VuRKD1_pro_:GUS* expression in the egg cell. C, *VuRKD1_pro_:GUS* expression in a sperm cell. D, *VuRKD1_pro_:GUS* expression in the vegetative cell. E, *VuGULL06_pro_:GUS* in the egg apparatus. F, *VuXTH32_pro_:GUS* expression in the vegetative cell. G, *VuGIM2_pro_:GUS* expression in the vegetative cell. Scale Bars: A, F, and G = 10 µm; B and D = 20 µm; C and D = 8 µm.

**Supplemental Figure. 7.**
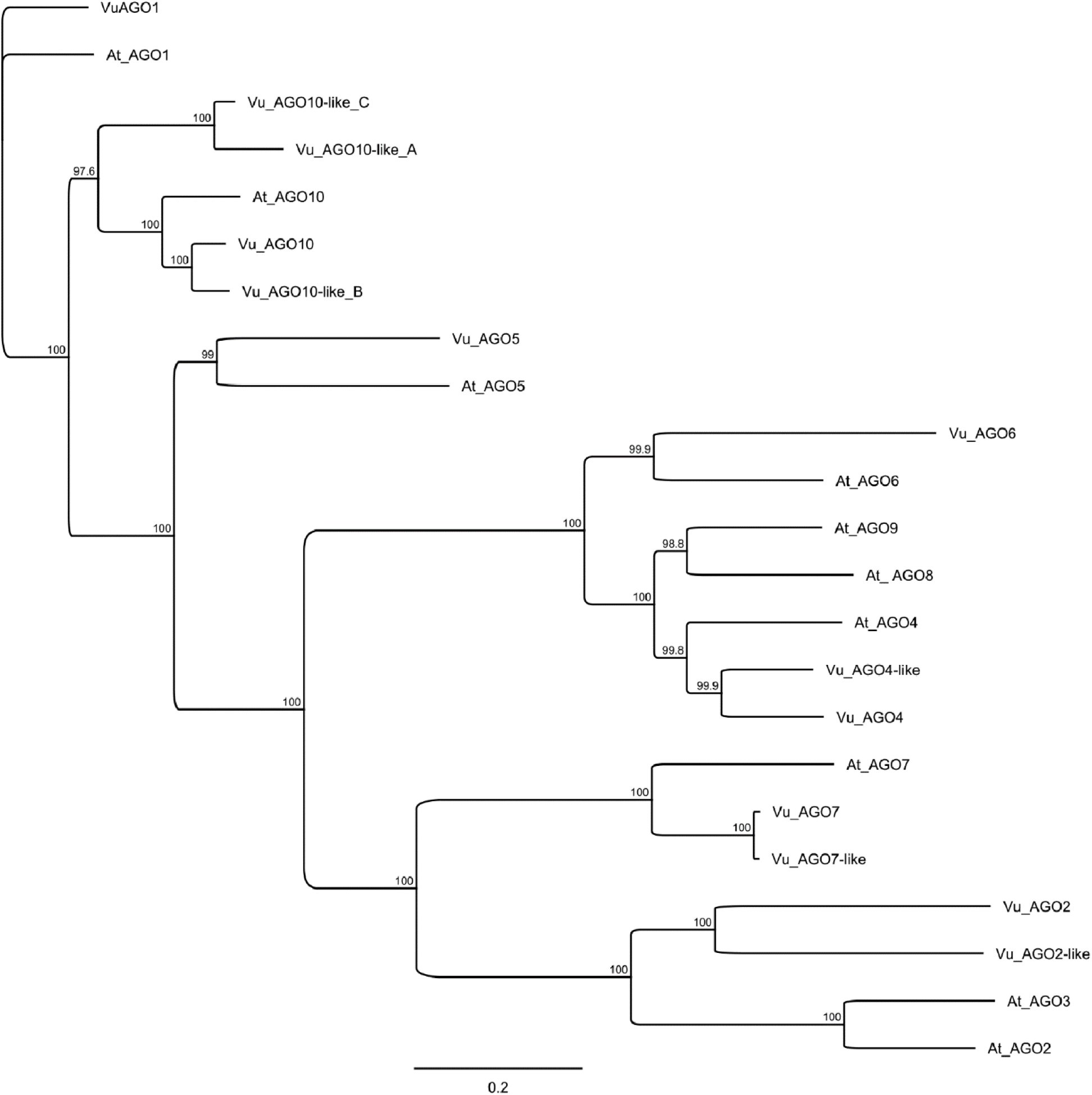
Phylogeny of the AGO family in cowpea and *Arabidopsis* A Jukes-Cantor Neighbour-Joining consensus tree showing the relationship between the AGO genes identified in cowpea (Vu) and those known in *Arabidopsis* (At). Parameters for tree building were ClustalW MSA with BLOSUM cost matrix, Gap open cost 10, Gap extend cost 0.1. Jukes-Cantor Neighbour-Joining consensus tree, no outgroup, Bootstrap resampling with 1000 replicates, 50% support threshold.

**Supplemental Figure. 8.**
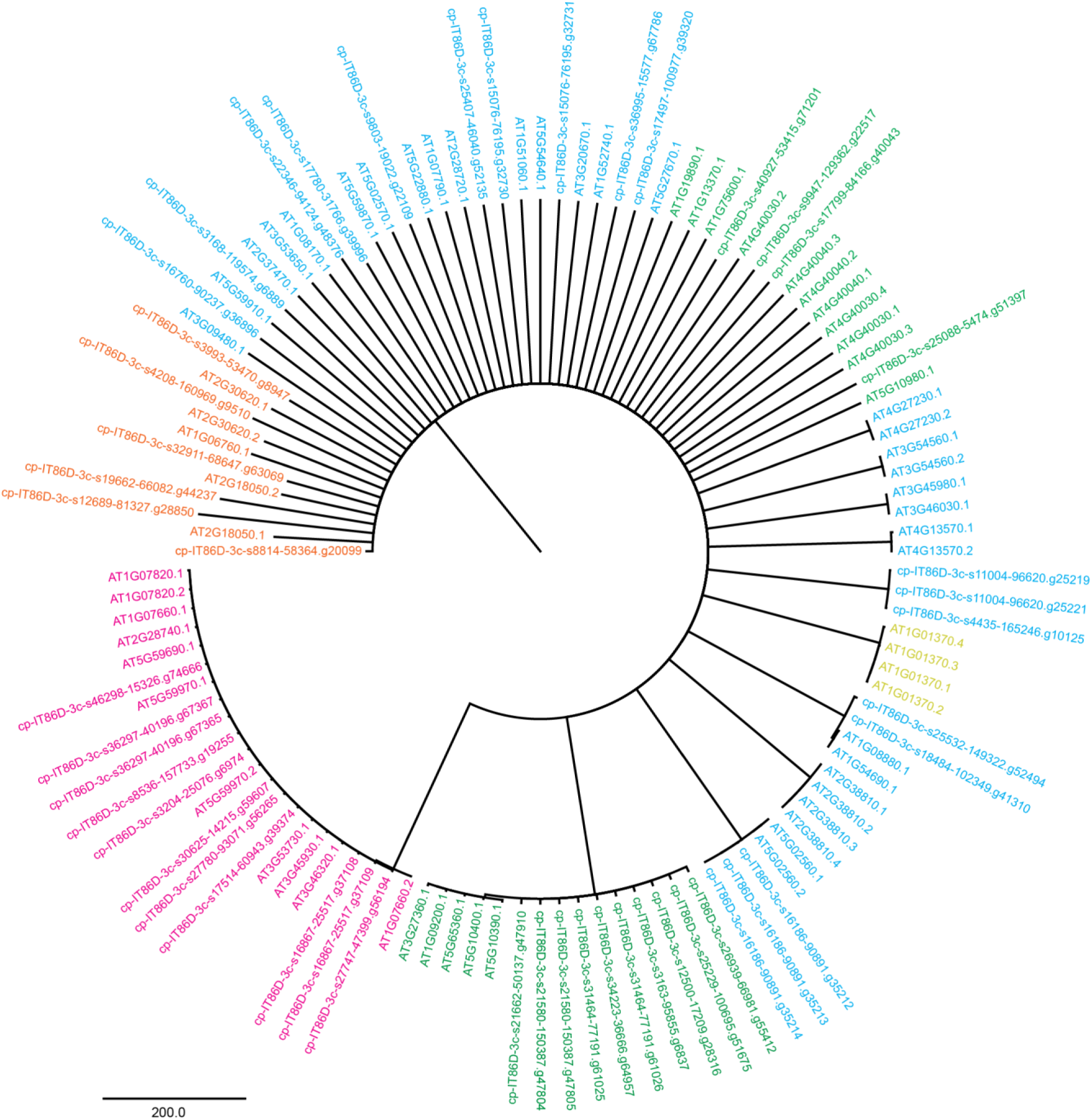
Phylogeny of *Arabidopsis* and cowpea histone proteins Jukes-Cantor Neighbour-Joining consensus tree showing the relationship between the *Arabidopsis* and cowpea histones is shown. Histone1 (orange), Histone2 (blue), Histone3 (green), Histone4 (pink), CenH3 (yellow). A Parameters for tree building were ClustalW MSA with BLOSUM cost matrix, Gap open cost 10, Gap extend cost 0.1. Jukes-Cantor Neighbour-Joining consensus tree, no outgroup, Bootstrap resampling with 1000 replicates, 50% support threshold.

